# A small molecule inhibitor of RNA-binding protein IGF2BP3 shows anti-leukemic activity

**DOI:** 10.1101/2025.04.14.648780

**Authors:** Amit K. Jaiswal, Georgia M. Scherer, Michelle L. Thaxton, Jacob P. Sorrentino, Constance Yuen, Milauni M. Mehta, Gunjan Sharma, Tasha L. Lin, Tiffany M. Tran, Amanda Cohen, Alexander J. Ritter, Rishi S. Kotecha, Jeremy R. Sanford, Robert D. Damoiseaux, Neil K. Garg, Dinesh S. Rao

## Abstract

The RNA-binding protein IGF2BP3 is an oncofetal protein overexpressed in B-acute lymphoblastic leukemia and is critical for leukemogenesis in experimental models. With cancer-specific expression, functional dispensability for normal development, and an unleveraged pro-oncogenic function in mRNA homeostasis, IGF2BP3 represents an excellent target. With no small molecule inhibitors of IGF2BP3 in clinical use, we undertook an effort to identify new IGF2BP3 inhibitors using biochemical methods. A biochemical screen, followed by a cell-based counter screen, led to the identification of compounds with protein-RNA interaction inhibition and leukemic cell growth-inhibitory activity. One of these compounds, designated I3IN-002, shows consistent cell growth-inhibitory activity, altered cell cycle and increased apoptosis in multiple leukemia cell lines, and is the most potent inhibitor of IGF2BP3 reported to date. I3IN-002 was tolerated in mice when administered intraperitoneally and showed potent anti-leukemic activity in a syngeneic transplantation model of MLL-Af4 leukemia. I3IN-002 inhibits the function of IGF2BP3, disrupting in situ binding of IGF2BP3 to target mRNAs, and altering IGF2BP3-dependent gene expression regulation. Furthermore, cell-free and cellular thermal shift assays as well as drug affinity responsive target stability assays support on target activity of I3IN-002 for IGF2BP3. Thus, the identification of I3IN-002 paves the way for the discovery of potent and selective small molecule inhibitors of IGF2BP3.

## INTRODUCTION

RNA binding proteins (RBPs) are important players in post-transcriptional gene regulation, and many show cancer-specific expression and may regulate oncogenesis via diverse mechanisms [1-5]. The RNA binding protein IGF2BP3 was initially discovered as a factor overexpressed in pancreatic cancer and one of three homologous proteins (IGF2BP1-3) that bound to the 5’-untranslated region of the IGF2 mRNA ^1^. Since its original description, a large body of work has congruently demonstrated its role as an oncogenic protein in a number of different cancers, and that IGF2BP3 binds to the 3’UTR of mRNA ^2^. In acute leukemia, we and others initially identified IGF2BP3 as an upregulated RBP in MLL-translocated B-ALL ^3, 4^. IGF2BP3 is transcriptionally induced by MLL-AF4, and overexpression of IGF2BP3 in bone marrow led to hyperproliferation in hematopoietic stem and progenitor cells in mice ^5^. Moreover, deletion of *Igf2bp3* in vivo significantly limited leukemia development and increased leukemia-free survival ^4, 6^. At baseline, the *Igf2bp3* knockout mouse did not show any overt phenotypes, suggesting that it is dispensable for normal development and homeostasis in the adult. Together these studies established IGF2BP3 as an oncogenic target in acute leukemia.

To investigate the molecular mechanisms of IGF2BP3’s action, we previously undertook iCLIP-seq, a technique designed to globally detect RNA molecules bound by IGF2BP3 in conjunction with differential expression analyses in cells that were depleted for IGF2BP3. These analyses revealed that IGF2BP3 most commonly binds to the 3’UTR of mRNA transcripts. The regulated mRNA transcripts included numerous downregulated oncogenes such as *HOXA9*, *CDK6* and *MYC* ^4, 7^. Determinants of IGF2BP3-RNA binding include not only primary sequence but also epitranscriptomic modification of target mRNAs with N6-methyladenosine (m^6^A) and spacing between short recognition sequences^6, 8–10^. Importantly, deletion of the RNA-binding domains of IGF2BP3 abrogated stabilization of these transcripts, and reversed IGF2BP3-driven pre-leukemic disease in vivo ^11^. Together these data suggest that IGF2BP3’s binding to specific mRNAs is central to its role in promoting oncogenesis in the hematopoietic system.

RBPs were previously under-represented in targeted therapy approaches due to their presumed “undruggable” status, recent efforts have uncovered small molecules targeting RBPs, including the IGF2BP family ^4, 12^. With the oncogenic role of IGF2BP3 established by not only our studies but also numerous studies in the literature, we sought to identify a small molecule inhibitor of the protein. To this end, we developed and validated a time-resolved Förster resonance energy transfer (TR-FRET) assay, using purified IGF2BP3 protein and a synthetic m^6^A-modified target RNA molecule to measure binding. A library of small molecule compounds was applied to this assay, with resultant hit compounds being re-confirmed in TR-FRET assays and counter screened in cell-based assays. Confirmatory studies to demonstrate activity in several leukemia cell lines with IGF2BP3 overexpression showed inhibition of cell growth and cell cycle progression, and increased apoptosis. The first of these compounds, referred to here as I3IN-002, shows activity against leukemic initiating cells in vitro, and has anti-leukemic activity in vivo. I3IN-002 directly interferes with the function of IGF2BP3, as assessed by gene expression analyses and RNA immunoprecipitation experiments that measure IGF2BP3-dependent gene expression and IGF2BP3-mRNA binding. Furthermore, cellular thermal shift assays, cell-free thermal shift assay (TSA), drug affinity responsive target stability (DARTS) support on-target activity for I3IN-002. I3IN-002 may prove useful as a tool compound in future biochemical studies involving IGF2BP3 and may also enable the development of a novel therapeutics for the treatment of leukemia and other hematologic cancers.

## METHODS

### A complete description of methods can be found in Supplementary Information. Cell lines and cell culture

All cell lines were maintained in standard condition in incubator at 37 °C and 5% CO_2_. Human B-ALL cell lines, RS4;11 (ATCC CRL-1873), NALM6 (ATCC CRL-3273), SEM (DMZ-ACC 546), PER785, REH (ATCC CRL-8286), and KASUMI-2 (DSMZ ACC 526), and immortalized MLL-Af4 transformed murine HSPCs were cultured as previously described ^6^. A full listing of reagents is provided in **Supp. Table 1**.

### Antibodies

Antibodies were used for Western blotting, FACS, and immunoprecipitation experiments as described ^4, 13, 14^. A list of antibodies is provided in **Supp. Table 2**.

### Protein Purification and High throughput assays

IGF2BP3-GFP-Flag over-expression plasmid was constructed by fusing full length IGF2BP3 CDS with GFP CDS and Flag CDS and cloned in pCDNA 3.0 vector. Protein was purified using standard methods and anti-Flag based affinity purification. Purified protein was added to pre-incubated Streptavidin-terbium plus biotinylated m6A-labeled RNA oligonucleotide (**Supp. Table 4**). The plate was incubated for 1 hr. at RT for binding and read on EnVision microplate reader (Perkin Elmer) with a dual PMT configuration using a 340 nm excitation and a dual emission mirror block with using a 495 nm emission filter for Terbium fluorescence (W1 channel) and a 525 nm emission filter for GFP fluorescence (W2 channel). The assay was miniaturized to 10uL in 384-well plates and an in-house compound deck at UCLA MSSR was applied to the assay. Data analysis was performed as described in Supplementary Methods.

### Counter-Screen and Cell-based Assays

Wild type and IGF2BP3-deleted SEM cell lines ^6^ were treated with compound, grown for 4 days under normal growth conditions, and assayed for cell growth in 384-well plates with CellTiter-Glo, a luminescence-based reagent described previously ^6, 13, 14^. Following identification of lead compounds, further measurements of apoptosis, cell cycle and proliferation were performed as we have previously described^6, 13, 14^.

### Compound Synthesis

Briefly, organic synthesis was performed based on utilizing commercially available fragments and a custom in-house strategy allowing for the design of the original compound and future analogs. Full description is provided in the Supplementary Methods.

### Assays to define on-target activity

A cellular thermal shift assay was performed on SEM cells treated with the test compound, based on published protocols with minor modifications described in the Supplemental methods ^14^. Thermal shift assay with purified protein, and Drug Affinity responsive target stability assays were performed as previously described ^15, 16^. Lastly, RNA immunoprecipitation and RNA-sequencing were performed using standard methods^4, 17^. RNA sequencing data will be deposited to the NCBI Short read archive with accession number XXXXXXX upon acceptance of the manuscript.

### Animal Experiments

For in vivo studies, C57BL/6J, B6.SJL-*Ptprc^a^ Pepc^b,^*/BoyJ (B6 CD45.1), and B6J.129(Cg)-Gt(ROSA)26Sor^tm1.1(CAG-cas9,-EGFP)Fezh/J^ (Cas9-GFP, BL/6J) were procured from The Jackson Laboratory. Preliminary toxicity studies were carried out to identify issues with small molecule delivery in non-transplanted mice. Primary murine leukemia cells (WT and Igf2bp3 KO) as previously described ^14^ were transplanted into busulfan-conditioned recipients. I3IN-002 was delivered intraperitoneally, 3 times a week for three weeks. Once the peripheral blood engraftment reached >20% at 4 weeks, the experiment was terminated and tissues were harvested to be analyzed by FACS, histology and RT-qPCR. A separate experiment examined overall and leukemia free survival. All of the animal experiments received Institutional Animal Research Committee approval at UCLA.

## RESULTS

### Disruptors of protein-RNA interactions are identified via biochemical screening

To identify putative inhibitors of IGF2BP3 (I3), we designed and validated a time-resolved Förster resonance energy transfer (TR-FRET) assay, based on binding of a GFP-tagged I3 (I3-GFP) to a biotinylated RNA oligonucleotide carrying a m^6^A modification (**Fig. 1a**). Small molecules that interfere with this interaction will lead to a loss of signal transfer, and subsequently decreased GFP fluorescence. For this assay, we generated a carboxy-terminal GFP- and FLAG-tagged IGF2BP3 protein-encoding mammalian expression vector, cloned into pcDNA3.1 (**Supp.** Fig. 1a). We confirmed a functional fusion protein based on bright fluorescence following transfection and recognition with anti-IGF2BP3 antibodies (**Supp.** Fig. 1b). The protein was purified by elution from fractions 3-11 on an anti-FLAG column (**Supp.** Fig. 1c) and subsequently concentrated and quantitated by Coomassie staining (**Supp.** Fig. 1d). Functionality of the protein was confirmed by RNA immunoprecipitation of target mRNAs following incubation of purified protein with cell extracts and pulldown with anti-Flag (**Supp.** Fig. 1e-g). Following purification and confirmation of RNA-binding activity, we confirmed that GFP fluorescence in the TR-FRET assay required the presence of all components (i.e., I3-GFP, biotinylated mRNA, and Terbium, **Fig. 1b**). The assay demonstrated saturation of the GFP fluorescence signal at approximately 50 nM of I3-GFP-F and 25 nM RNA oligonucleotide (**Fig. 1c and Supp.** Fig. 1h). Next, we evaluated assay reproducibility across multiple experiments performed across multiple days, finding excellent concordance in GFP fluorescence and Z’-values (**Supp.** Fig. 1i-j).

**Figure 1.**
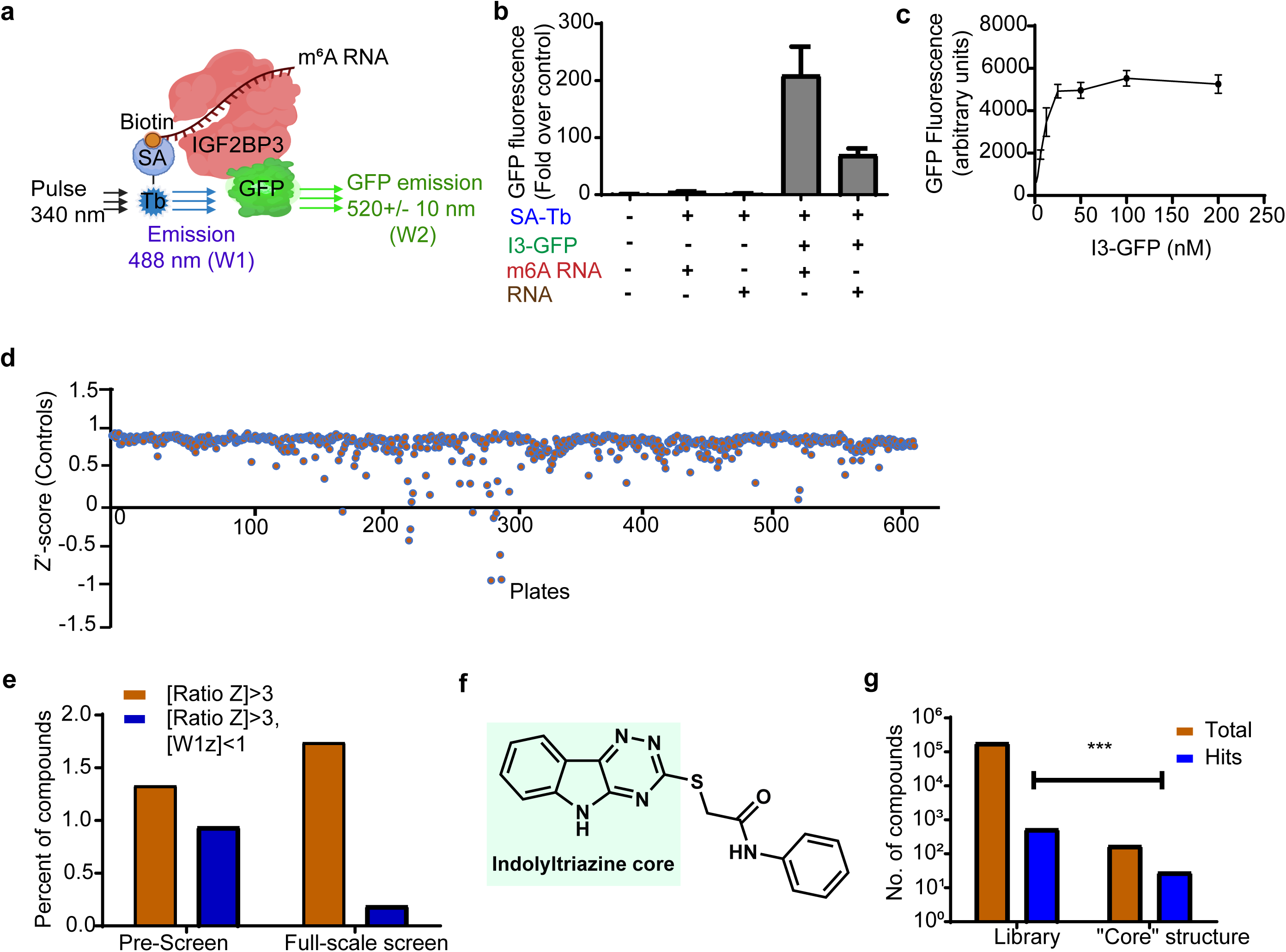
Design and execution of a high throughput assay to identify small molecule inhibitors of IGF2BP3. a. Schematic of assay. IGF2BP3-GFP fusion protein is bound to an m^6^A-modified, biotinylated RNA oligonucleotide, which in turn is bound to Streptavidin Terbium (SA-Tb). Förster resonance energy transfer occurs when the protein and RNA are bound. b. GFP Fluorescence (fold over control) plotted for different combinations of Streptavidin Terbium (SA-Tb), IGF2BP3-GFP (I3-GFP), m6A-modified RNA (m6A RNA) or unmethylated, biotinylated RNA (RNA). c. GFP Fluorescence (arbitrary units) as a function of IGF2BP3-GFP protein concentration. d. High throughput screening assay Z’-scores for control reactions across 600+ plates screening over 186,978 compounds. e. Screening was conducted first with a limited set of compounds (preliminary screen) prior to a full-scale screen. Fluorescence ratios were calculated as W1/W2 where W1 is the Terbium fluorescence and W2 is the GFP fluorescence. Shown are the percent of compounds satisfying a single criterion (i.e., Ratio Z) or two criteria (Ratio Z and W1 Z). f. First “core structure” identified from the screen. The indolyltriazine substructure was conserved across multiple hits from screening in d and e. g. Comparison of total number of hits in the library, versus hits amongst compounds containing the first core structure; Fisher’s exact test, p<0.0001.

To adapt the assay to high throughput screening, we first successfully miniaturized the assay to 10 μL, while maintaining σ+ <20% and Z’>0.5, with highly reproducible GFP fluorescence (**Supp.** Fig. 1k). Following this validation, a high throughput screen of 196,567 compounds was completed in two stages, a preliminary screen, and a full-scale screen. We were able to successfully complete the screen: across 617 plates that were analyzed for this assay, we achieved >99% of plates that showed standard deviation <20% and 95% of plates showed a Z’ score greater than 0.5 (**Supp.** Fig. 2c and **Fig. 1d**). Plates that showed a low Z’-score were excluded from downstream hit confirmation. To define hits, we leveraged the preliminary screen to increase the specificity of hit definition (**Supp.** Fig. 2 **a-c**). Using only the primary criterion, defined as |Ratio Z|>3 (where Ratio is the ratio of terbium fluorescence/GFP fluorescence), we obtained a hit rate of 1.33% in the preliminary screen, and 1.73% in the full-scale screen (**Fig. 1e**). Using a secondary criterion, where Terbium fluorescence (“W1”) was within one standard deviation of the mean for the plate (i.e., |W1 Z|<1), this reduced the hit rate to 0.94% and 0.19% in preliminary and full-scale screens respectively (**Fig. 1e**). The reduced hit rate in the full-scale screen therefore is most likely due to increased variability in terbium fluorescence. Further refinements of hits, by an alternate secondary criterion definition (mean and standard deviation calculated for the experimental run, rather than the plate) and manual curation of the data when irregularities were observed, led to an initial, high-stringency list of 417 compounds that were further analyzed. Of the 417 hits out of 186,978 evaluable compounds, 29 contained a core structure with an indolyltriazine group (**Fig. 1f**). Remarkably, the library only contained 174 total compounds with the general structure depicted in **Fig. 1f**, representing ∼74-fold enrichment for these compounds, a highly statistically significant result (**Fig. 1g**; p<0.0001; Fisher’s exact test). Together, our results identify a number of small molecules with in vitro activity against the RNA binding activity of IGF2BP3.

### Cell-based counter screen identifies molecules with anti-leukemic activity

To further characterize the hits identified from the screen, we first performed a confirmatory TR-FRET assay, with two experiments consisting of three replicates each. Concordant TR-FRET findings were characterized as Tier 1, 2 and 3, when findings were reproduced in 2/2, 1/2, or 0/2 experiments respectively. 89.3% (371/417) of hit compounds and 93.1% (27/29) of indolyltriazine compounds were Tier 1 or Tier 2 hits, respectively (**Supp.** Fig. 2d). To further examine these confirmed hits, we turned to a cell-based assay (**Fig. 2a**). Here, we reasoned that a compound that was specific for IGF2BP3 would only show activity in SEM cell lines that contained intact, increased expression of the protein (SEM-WT), with reduced activity in cell lines with experimental deletion via CRISPR-Cas9 (SEM-I3KO) ^18^. Cells were treated with compound, grown for 4 days under normal growth conditions, and assayed for cell growth with CellTiter-Glo, a luminescence-based reagent described previously ^6, 13, 14^. Fold change from control (i.e., no compound treated wells) was plotted for SEM-WT versus SEM-I3KO. Remarkably, there were compounds that showed enhanced reduction of cell growth in SEM-WT, as opposed to SEM-I3KO (**Fig. 2b**). Using the cutoff (Growth Ratio<0.7 OR WT Growth<0.7), we identified 16 compounds. We then chose three promising compounds, designated **1**, **2** and **3** here, and tested them across a range of concentrations on SEM cells (**Fig. 2c-e**). Of these, **2** and **3**, which we later designated as **I3IN-002** and **I3IN-003**, showed differential activity in the two cell lines, with **2** showing a promising IC_50_ of ∼2 µM in wild-type SEM-cells (**Fig. 2d**).

**Figure 2.**
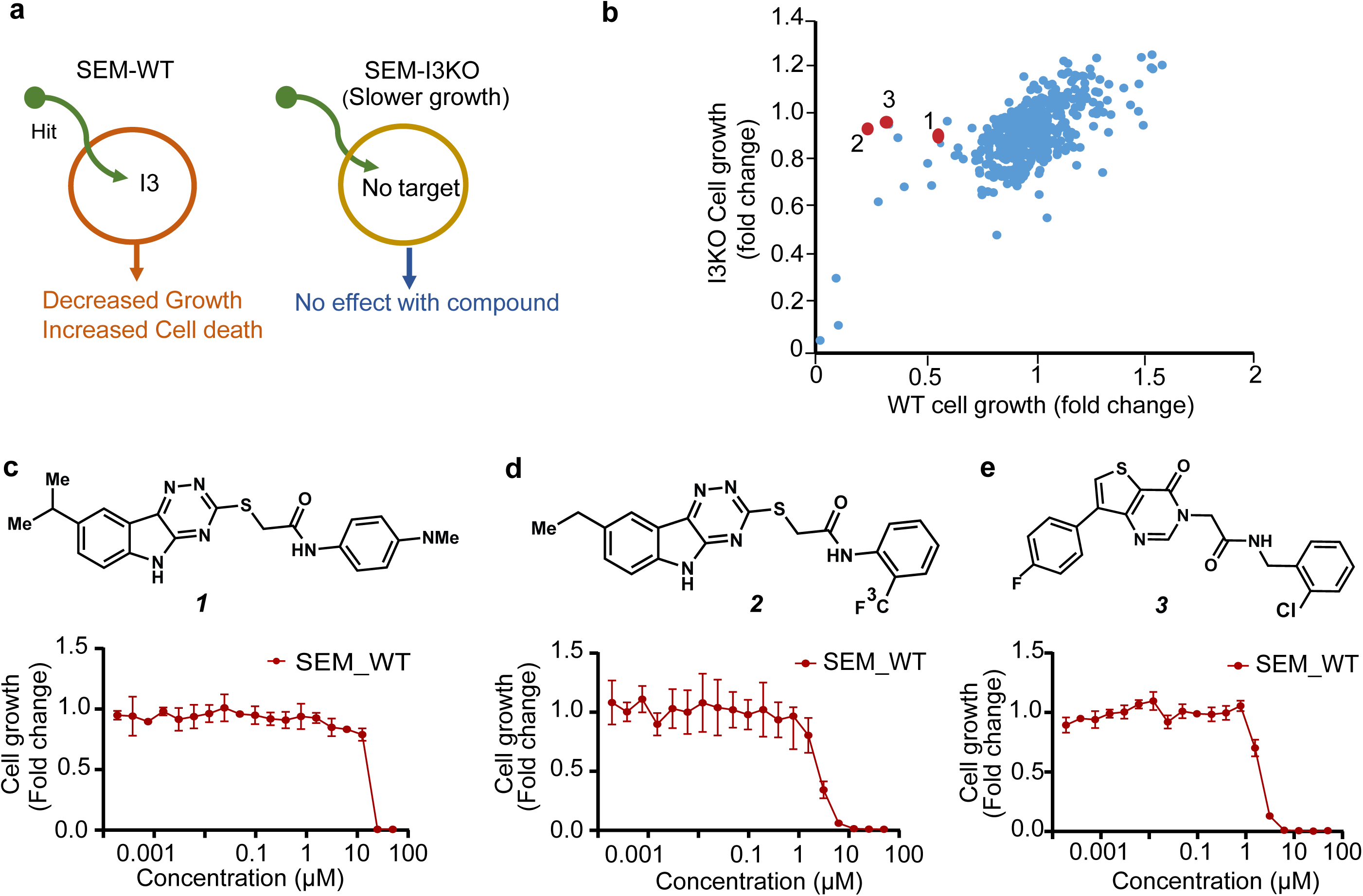
Identification of molecules with biochemical activity and IGF2BP3-dependent anti-proliferative properties. a. Counter-screen strategy: SEM cells with (SEM-WT) and without IGF2BP3 (SEM-I3KO) are treated in a medium-throughput format with the small molecules identified from the biochemical screen. b. Scatter plot examining the relationship between relative changes in cell growth, measured by CellTiterGlo and normalized to DMSO control, in SEM-WT (x-axis) and SEM-I3KO (y-axis). Compounds *1*, *2* and *3* (red-brown) were noted to have a preferential effect on SEM-WT and a lesser effect in SEM-I3KO. Cells were treated with 5 µM solutions of test compounds. c-e. Structures of compounds *1*, *2* and *3* (top) and corresponding concentration dependent changes in cell growth rate (bottom) for the three compounds with favorable characteristics on screening. Growth rate was measured using CellTiterGlo and normalized to DMSO control in SEM-WT cell line.

### I3IN-002 inhibits cell proliferation and cell cycle progression and promotes apoptosis

To enable further biological evaluation, we sought to prepare **I3IN-002** in high purity and in larger quantities compared to what was available commercially. Moreover, chemical characterization data for **I3IN-002** was not identified in the literature. To access **I3IN-002**, we performed the synthetic route shown in (**Fig. 3a**). First isatin **4** was treated with thiosemicarbazide (**5**) and potassium carbonate in methanol at ambient temperature to give hydrazone formation. The hydrazone intermediate was subjected to potassium carbonate in water at reflux to give triazinoindolothione **6** ^14^. In the next step, intermediate **6** was treated with alkyl chloride **7** in the presence of potassium carbonate in DMSO ^19^. This delivered **I3IN-002**, which was isolated as a powder. The purity of **I3IN-002** was deemed to be >97% by both quantitative ^1^H NMR and HPLC analysis.

**Figure 3.**
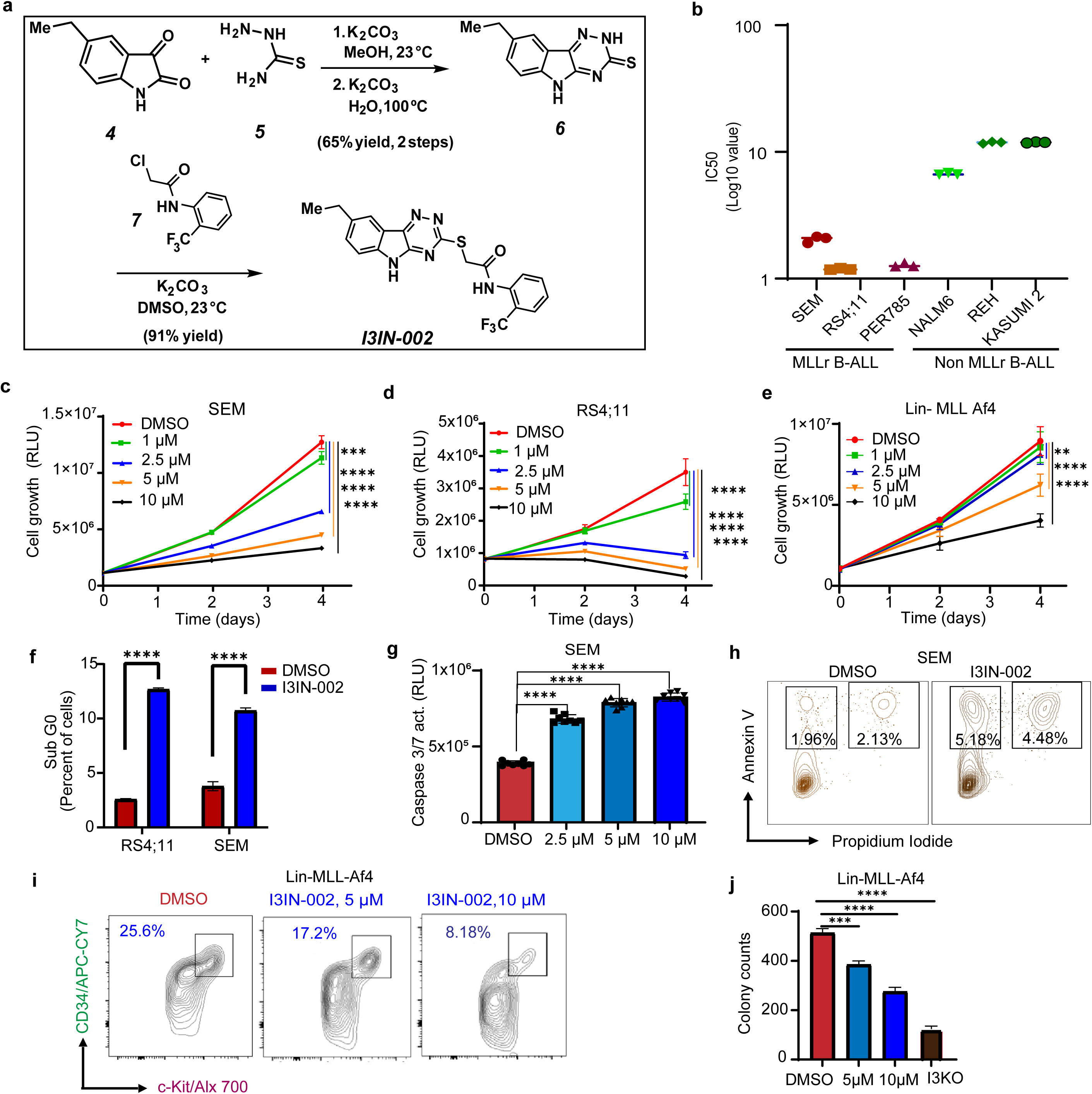
I3IN-002 alters cell growth, cell cycle, and apoptosis in human and mouse leukemic cells. a. Chemical synthesis of **I3IN-002**. b. IC50 values of I3IN-002 on MLLr B-ALL and B-ALL cell lines. c-e. Concentration-dependent alterations of cell growth curves over 4 days of cell culture with I3IN-002 treatment, assessed by CellTiterGlo assay, and plotted as relative luminescence units (RLU). SEM (c), RS4;11 (d), and Lin-/MLL-Af4 (e) cells were assayed in this experiment. Data represent mean and standard error of mean in 8 replicates. f. Measurement of SubG0 cells by propidium iodide staining in SEM and RS4;11 cells, respectively, following treatment with 5 µM I3IN-002 or DMSO. Data represents mean and standard error of mean of the calculated percentage of cells at each stage of cell cycle with 3 replicates. g. Measurement of apoptosis induction, by Caspase 3/7 activity assay in SEM cells. Data represents mean and standard error of mean with 8 replicates. h. Measurement of cell death by orthogonal method of Annexin V and propidium iodide staining in SEM cells. Shown are the % of cells that are single or double positive for propidium iodide and Annexin V. i. Quantitation of a leukemic stem-cell enriched population by assaying the CD34+ c-kit+ population following in vitro treatment of Lin-/MLL-Af4 cells in culture. j. Measurement of colony formation in methylcellulose following treatment of a murine model of MLL-Af4 driven leukemia ^13, 14^ with I3IN-002. Data represents the mean number of total colonies for each condition, with 3 replicates per condition. I3KO represents genetic knockout of IGF2BP3, which results in reduced colony formation For Figure 3, statistical significance was evaluated by student’s T-test, *p<0.05; **p<0.01; ***p<0.001; ****p<0.0001.

To characterize the activity of I3IN-002, we tested the compound in a number of leukemia models, including SEM, RS4;11, PER 785, KASUMI-2, NALM6, REH, and MLL-Af4 Lin-cells. IC50 experiments performed on these cell lines showed the compound is more potent in B-ALL cell lines with MLL-AF4 translocation, with lower IC50 values compared to other B-ALL cells (**Fig. 3b**). Treatment with a single dose of I3IN-002 resulted in a dose-dependent growth inhibition of SEM, RS4;11 and Lin-MLL-Af4 cells at the concentrations tested in our assays (**Fig. 3c-e**). Moreover, cell cycle profiling using propidium iodide in both SEM and RS4;11 cells showed an increase in sub-G0 cells (**Fig. 3f and Supp.** Fig. 3a-3c); changes in other phases of cell cycle were not as consistent. This may be in part, because in SEM cells the G2-M transition genes such as CDK1 and CCNB2, which are also IGF2BP3 targets, were downregulated due to compound treatment (**Supp.** Fig. 3d-3e). Next, we analyzed the effect of I3IN-002 on cellular apoptosis. These studies showed an increase in apoptosis, as detected by Caspase 3/7 activity in both SEM and RS4;11 cells (**Fig. 3g and Supp.** Fig. 3f, respectively). Treatment also led to a numerically small but statistically significant increase in Annexin V+ cells as well as in necrotic cells (Annexin V+ and Propidium iodide+) in both SEM and RS4;11 cells (**Fig. 3h and Supp.** Fig. 3g-i). Together, these studies indicate that I3IN-002 impacts cell growth, cell cycle and apoptosis in a variety of target cell lines, with an enhanced growth inhibitory effect in MLL-AF4-driven leukemia.

### Reduction of leukemia initiating cells and leukemia engraftment in vivo by I3IN-002

Given our previous work showing that genetic ablation of *Igf2bp3* led to a reduction in leukemia initiating cells, we next assayed whether I3IN-002 caused an effect on plating efficiency in methylcellulose colony formation assays. Like genetic deletion, I3IN-002 treatment led to a reduction in total colony counts in a dose-dependent manner and a reduction in the fraction of cells that showed expression of CD34 and c-kit (**Fig. 3i-j and Supp.** Fig. 3j). With these findings, we next evaluated the compounds in vivo.

For in vivo experiments, mice syngeneically transplanted with the Lin-MLL-Af4 cells (either WT or I3KO, **Supp.** Fig. 4a) ^20^ were injected intraperitoneally with I3IN-002 or vehicle control using the schedule indicated (**Fig. 4a**). This model demonstrates a rapid and lethal leukemia that primarily involves the bone marrow and spleen, with lesser involvement of the peripheral blood. Here, we used the genetic deletion of *Igf2bp3* as a positive control for the predicted effect of IGF2BP3 inhibition, as genetic ablation prevents leukemia development(14,15). Upon harvest at 4 weeks post-transplant, WT-transplanted mice that received I3IN-002 showed significantly reduced spleen size and weight compared to vehicle treated control mice, but larger than I3KO-transplanted mice (**Fig. 4b and 4d**). Histologically, we observed lower levels of leukemic engraftment in the liver (compare cellular area next to arrow) and reduced architectural distortion and lower leukemic engraftment in the spleen (compare size of dotted areas and intervening cellularity) in I3IN-002 treated mice, and as expected in I3KO (**Fig. 4c**). Similar reductions in engraftment were also noted in lung and kidney (**Supp.** Fig. 4f). FACS analysis revealed that in vivo treatment reduced the number of CD34+ c-kit+ cells in both the spleen and the bone marrow (**Fig. 4e (spleen), 4h and 4k**). In the spleen, leukemic engraftment and the proportion of CD11b+ cells and of CD34+ Sca1-cells were reduced significantly (**Fig. 4f-g** and **Supp.** Fig. 4b-4c). In the bone marrow, overall leukemic engraftment was not different following I3IN-002 treatment compared to vehicle, but significant reductions in CD11b+ and CD34+ Sca1-cells were noted (**Fig. 4j** and **Supp. 4d-4e**). Further, we found significant improvement in overall survival following I3IN-002 treatment in a separate set of endpoint experiments (**Fig. 4l**). Together, these data indicate a significant effect of I3IN-002 on leukemic burden, corresponding with a reduction in the leukemic initiating cell fraction, thus inhibiting leukemogenesis and extending survival.

**Figure 4.**
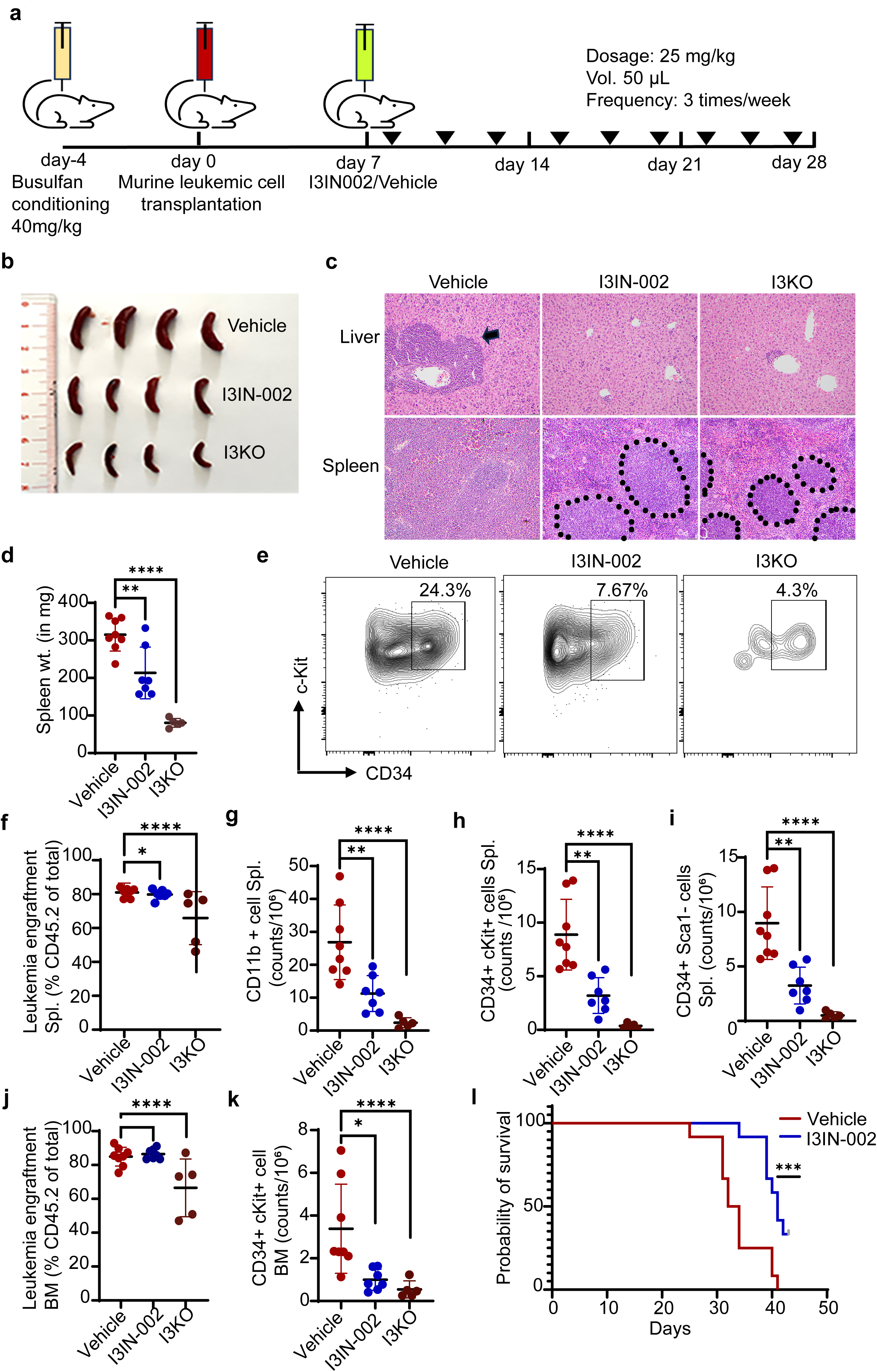
I3IN-002 inhibits leukemic growth in vivo. a. Schematic of mouse experiment. b. Gross pathology photographs of murine spleens from mice syngeneically transplanted with Lin-/MLL-Af4 cells. I3KO are genetic knockouts of IGF2BP3, which markedly reduced leukemic engraftment. c. Histology of liver (top) and spleen (bottom) from treated mice, as indicated. Arrow indicates a leukemic infiltrate in the liver in the portal area. Dotted lines indicate splenic white pulp, with leukemic infiltrates in the red pulp (area outside the dotted lines). Hematoxylin and eosin staining, original magnification, 200X. d. Splenic weights from treated mice, as indicated. e. FACS staining for CD34 and c-kit in the spleen of treated mice, as indicated. f-i. Quantification of overall leukemia engraftment and various leukemia subsets in the spleen. j-k. Quantification of overall leukemia engraftment and various leukemia subsets in the bone marrow. l. Kaplan-Meier survival curve shows effect of I3IN-002 injection on overall survival in syngeneic bone marrow transplant experiment. Data in e-k represents one experiment composed of the following numbers of animals: Vehicle, n=8; I3IN-002, n=7; I3KO, n=5. Statistical significance was evaluated by student’s T-test, *p<0.05; **p<0.01; ***p<0.001; ****p<0.0001.

### I3IN-002 inhibits the molecular function of IGF2BP3 in post-transcriptional gene regulation

Prior work by our group and others ^14^ has shown that IGF2BP3 functions in post-transcriptional gene regulation by binding to mRNA molecules and regulating their expression. To test how I3IN-002 impacts IGF2BP3 function, we first performed RNA immunoprecipitation (RIP) following treatment with I3IN-002. We found that four mRNAs that we have previously found to be associated with IGF2BP3, *CDK6, MYC, BCL2,* and *HOXA9*, were decreased in the immunoprecipitate following IGF2BP3 pulldown (**Fig. 5a-b** and **Supp.** Fig. 5a-b). Next, to assess *in situ* binding of I3IN-002 to IGF2BP3, we performed a cellular thermal shift assay, finding a difference in thermal stability, as assessed by Western blot (**Fig. 5c-d**). Direct interaction of I3IN-002 with IGF2BP3 was confirmed using orthogonal methods. Thermal shift assay (TSA) using purified full length IGF2BP3 demonstrated stabilization of IGF2BP3 with compound (**Fig. 5e**), while DARTS demonstrated I3IN-002 dependent inhibition of protease mediated degradation of IGF2BP3 (**Supp.** Fig. 5c). To understand the downstream consequences of IGF2BP3 inhibition, we performed RNA-seq of DMSO-treated, I3IN-002-treated and I3KO SEM cells. These studies resulted in the identification of differentially expressed genes between SEM-WT and SEM-I3KO (n=319 genes) as well as between SEM-WT and SEM-WT-I3IN-002 treated cells (n=226 genes). Volcano plots demonstrated that the overall changes in gene expression were more pronounced in the genetic knockout of IGF2BP3 compared to I3IN-002 treated cells (**Fig. 5f**). Remarkably, 63 genes were common between the two datasets, which represents a highly significant overlap (nearly 28% of genes regulated by I3IN-002 treatment; Hypergeometric test, p-value< 10^-16^; **Fig. 5g**). Gene set enrichment analyses of the overlap genes between I3KO and I3IN-002 treated cells showed many pathways that we have previously associated with IGF2BP3 function, including response to oxidative stress, cell activation, cellular response to cytokine stimulus and neutrophil degranulation (**Fig. 5h**). A selected panel of differentially regulated genes were confirmed by RT-qPCR (**Fig. 5i-n and Supp.** Fig. 5e-f), and Western blotting confirmed the downregulation of CDK6 and BCL2 proteins that we have previously observed with I3KO (**Fig. 5o**). Paralleling the findings in cell culture, several IGF2BP3-regulated mRNAs, including Myc, Cdk6, and Hoxa9 were downregulated in spleen samples from mice treated with I3IN-002 based on RT-qPCR (**Supp.** Fig. 5g-5l). Together, these findings support a mechanism wherein I3IN-002 is impacting the function of IGF2BP3 in its critical role as a post-transcriptional regulator of mRNA homeostasis.

**Figure 5.**
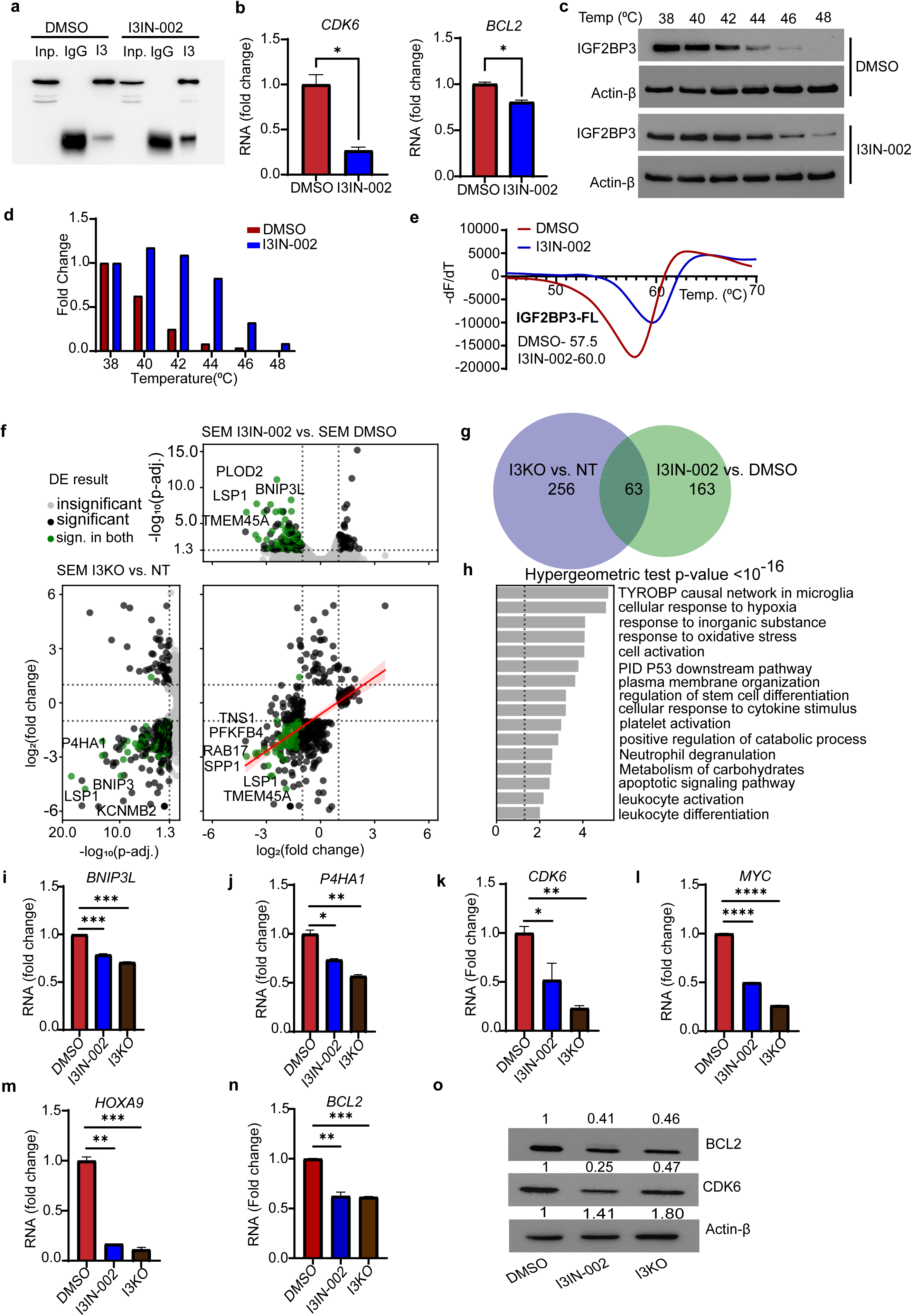
I3IN-002 shows on-target activity in leukemia cells. a. Western blot analysis of RNA immunoprecipitation samples following treatment of SEM cells with 5 µM I3IN-002. Input represents the input cell lysate; IgG represents a control immunoprecipitation with isotype control: and “IP” represents anti-IGF2BP3 immunoprecipitation. b. Quantification of *CDK6* and *BCL2* RNA recovered from RNA immunoprecipitation experiments with or without treatment with I3IN-002. c-d. Cellular thermal shift assay; Western blots of IGF2BP3 and beta-actin are shown following treatment of cells with I3IN-002 or DMSO control, followed by heat-mediated denaturation and removal of denatured protein (c) and quantification of western blot (d). e. Thermal shift assay using purified full length IGF2BP3 protein. f. Volcano plots of differentially expressed genes with genetic IGF2BP3 knockout (left) and with I3IN-002 treatment (top) in SEM cells, using DESeq analysis on RNA-sequencing data from SEM cells treated with DMSO, I3IN-002 and SEM/I3KO. g. Venn diagram of shared differentially expressed genes with IGF2BP3 knockout and I3IN-002 treatment in SEM cells. Hypergeometric test of overlap showed a p-value<10^-16^. h. GSEA analysis of the overlapped genes between I3KO and I3IN-002 treated cells. i-n. RT-qPCR based confirmation of genes showing common differential regulation between I3IN-002 treatment and genetic knockout of IGF2BP3 (I3KO). Data are shown as fold change, following normalization to DMSO, with 3 replicate measurements per sample. o. Western blot measurement of CDK6, and BCL2, which are known targets of IGF2BP3.

## DISCUSSION

The RNA-binding protein IGF2BP3 is a potent oncofetal protein, as demonstrated by our work and that of many others. In this manuscript, we describe the successful completion of a high-throughput biochemical screening assay that utilizes the RNA-binding function of IGF2BP3 as the discriminant for small molecule activity. Combining the biochemical screen with a cell-based assay to detect IGF2BP3-dependent effects on leukemic cell growth and survival, we identified 16 compounds that are likely to have activity in an IGF2BP3-specific manner. The first of these compounds to be confirmed is I3IN-002, which shows an IC_50_ of approximately 2 μM in initial assays. Specifically, I3IN-002 inhibited cell growth, impacted cell cycle progression of treated cells, and increased apoptosis. I3IN-002 inhibited leukemia initiating cells and showed in vivo anti-leukemic activity. Furthermore, I3IN-002 impacted global gene expression in a manner similar to genetic deletion of IGF2BP3. CETSA, TSA, DARTS and RNA immunoprecipitation experiments supported on-target activity against IGF2BP3.

RNA binding proteins have recently been appreciated as biologically and clinically significant regulators of gene expression ^12, 21–23^. This recognition has led to a growing effort to target RNA binding proteins, but challenges exist ^12, 24^. First, many cancer-associated RNA binding proteins are also required for normal homeostasis, and hence, targeting these proteins might engender significant toxicity. The IGF2BP family of proteins have emerged as cancer-specific or cancer-enriched proteins, and IGF2BP3 is an oncofetal protein, making this an attractive cancer target. Second, while oligonucleotide-based therapies hold great promise, targeting of many tissues, including cancer, remains a major challenge in the field. Hence, small molecule targeting holds many advantages, including delivery, with development of reproducible and straightforward synthesis methods allowing for future large-scale manufacturing and distribution. Our efforts here have resulted in the some of the first small molecules identified by empirical, experimental approaches to target IGF2BP3.

In recent years, several small molecules have been reported to inhibit the IGF2BP family of proteins. Regulators of expression include an isocorydine derivative, 8-amino-ICD, which was reported to target hepatocellular carcinoma and decrease IGF2BP3 expression at EC_50_ of 20 μM or greater ^25^. In non-small cell lung cancer, isoliquiritigenin is also able to down-regulate IGF2BP3 expression in a dose-dependent manner ^26^. Most recently, a virtual screening approach identified AE-848 as an inhibitor of IGF2BP3 with an IC_50_ of ∼10-40 μM in ovarian cancer ^27^. Other IGF2BP proteins have been reported to be targeted by small molecules: BTYNB targets IGF2BP1 (IC_50_ of 5-10 μM); JX1 targets IGF2BP2 (IC_50_ of 10-20 μM) in acute T-lymphoblastic leukemia, and CWI1-2 targets IGF2BP2 also (sub-micromolar IC_50_ in AML cell lines)^18, 28, 29^. Our discoveries here indicate that I3IN-002 is the most potent inhibitor of IGF2BP3 reported to date, and the first in hematologic malignancy, providing the impetus for further drug discovery efforts.

Biologically active small molecules containing indolyltriazine cores possess potent and selective activity across a variety of indications. Other anti-cancer compounds containing the motif include an inhibitor of tyrosine phosphatase SHP-2^30^, and a commercially available tool compound, Inauhzin, which acts as a SIRT-1 inhibitor ^31^. Further, this motif has been also found in antibiotics, α-glucosidase inhibitors, and targeted p-glycoprotein inhibitors, amongst other classes of biologically relevant molecules ^32–35^. Notably, many of these compounds are demonstrated to be highly selective. This relatively diverse set of targets is consistent with our findings that small changes in the structure can markedly alter the bioactivity of the indolyltriazine in acute leukemia (i.e., only 2 out of 29 TR-FRET-active indolyltriazine compounds tested showed bioactivity). In other words, the specificity of the compounds may be highly malleable with chemical modifications of the side chains. Our initial efforts indicate that I3IN-002 can be readily synthesized using widely commercially available component molecules. This information should enable future efforts to develop compound analogs of I3IN-002 that display selective binding to IGF2BP3, without off-target effects. Importantly, while the indolyltriazine is seemingly privileged with respect to bioactivity, it is not found in any commercially available drugs to date.

It is also of interest to note that we recently showed that IGF2BP3 genetic ablation was additive with menin inhibition in MLL-AF4 leukemia ^36^. While outside the scope of this report, compounds generated here could therefore be useful in combinatorial therapeutic approaches with menin inhibition, inhibiting leukemia at the transcriptional and post-transcriptional levels. In addition, IGF2BP3 targets include a number of key oncogenic proteins, which are themselves the targets of advanced small molecule development programs. For example, CDK6 is a direct target of IGF2BP3 that is upregulated by the latter’s activity, and anti-CDK4/6 inhibitors have been developed as potent agents in cancer therapy, for example, in breast cancer ^14^. BCL2 is also a direct target of IGF2BP3, and the BCL2 inhibitor Venetoclax has shown activity in a number of hematologic malignancies, including B-ALL and AML. Other IGF2BP3 targets, such as MYC, which have long been elusive as drug targets, are also now being actively pursued with small molecule approaches ^37^. Hence, our development of IGF2BP3 small molecule inhibitors could be especially important in future combinatorial therapy approaches, and is an important future direction for our work.

While this work contains important advances, showing that disrupting IGF2BP3-RNA binding can be the basis of a therapeutic approach in selected hematologic malignancies, we acknowledge that there are areas for important future work. In vitro studies of binding affinities have yet to be completed, due to difficulties in purifying full-length protein in quantities sufficient for detailed biochemical analyses. We believe this is because the protein may be prone to liquid-liquid phase separation, and prior work has focused on individual domains within the protein, yielding crystal structures of protein subdomains only ^38–41^. Moreover, pharmacological parameters of this compound/derivatives will require optimization. Thus, future directions will involve analog synthesis and biochemical evaluation to ultimately improve and validate selective binding to IGF2BP3, while also optimizing drug-like properties.

Overall, our study has led to the discovery and validation of I3IN-002, a small molecule that inhibits IGF2BP3. While our work has thus far focused on the role of IGF2BP3 in acute leukemia, it is known that IGF2BP3 is overexpressed in a range of B-cell malignancies, including Diffuse Large B-cell lymphoma, Burkitt lymphoma and others^42–47^. Moreover, it is estimated that up to 15% of all human cancers overexpress IGF2BP3, including both hematologic and non-hematologic malignancies, with high levels correlating with aggressive tumor behavior by expression and functional analyses ^3, 21, 48–53^. Consistent with the notion of being an oncofetal protein^54^, our work has determined that IGF2BP3 appears to be largely dispensable in homeostatic development ^55^, providing a significant therapeutic window for targeting. It should be noted that IGF2BP3 expression is quite variable within different cancer types, and hence, future indications would need to be carefully defined. We speculate that IGF2BP3 expression could be evaluated in different cancer types by widely available techniques such as immunohistochemistry; and compounds such as I3IN-002 or future derivatives could be administered as a precision medicine intervention in carefully selected patients. Hence, I3IN-002 serves as a promising lead compound for future drug discovery efforts aimed at identifying selective inhibitors of IGF2BP3, particularly for the treatment of acute leukemia.

## Supporting information

All Supplemental Information

## ACKNOWLEDGMENTS

We thank Dr. Stuart Conway and Keefe Oei for helpful conversations, advice and protocols for biophysical assays. We thank Jaspal Bassi and Dr. John Colicelli, for helpful discussions regarding the research. This work was supported by CIRM DISC-2-13456 from the California Institute of Regenerative Medicine (DSR), R01CA264986 from the National Institutes of Health (DSR, JRS), R03CA251854 (DSR), the Jonsson Comprehensive Cancer Center (JCCC) (DSR), the Gary & Barbara Luboff Mitzvah Fund (DSR) and the UCLA Innovation Fund Award (DSR). Flow cytometry was performed in the UCLA JCCC and Center for AIDS Research Flow Cytometry Core Facility that is supported by National Institutes of Health awards AI28697, and award number P30CA016042, the JCCC, the UCLA AIDS Institute, and the David Geffen School of Medicine at UCLA. We are especially indebted to the Molecular Screening Shared Resource team for setup and execution of small molecule related screening assays. The MSSR is also supported by the Jonsson Comprehensive Cancer Center (NIH award number P30CA016042, Cancer Center Support Grant). These studies were also supported by the National Science Foundation (DGE-2034835 for GMS and DGE-1650604 for MMM), shared instrumentation grants from the NSF (CHE-1048804), the National Center for Research Resources (S10RR025631), and the NIH Office of Research Infrastructure Programs (S10OD028644).

## AUTHOR CONTRIBUTIONS

A.K.J.: Designed research, Performed experiments, Acquired data, Analyzed data, Generated Figures, Wrote manuscript, Experimental Lead

G.M.S.: Performed experiments, Acquired data, Analyzed data, Assisted in generating figures M.L.T.: Performed experiments, Acquired data, Analyzed data

J.P.S.: Performed experiments, Acquired data, Analyzed data C.Y.: Performed experiments

M.M.M.: Performed experiments, Acquired data, Analyzed data G.S.: Performed experiments

T.M.T.: Performed experiments

T.L.L.: Performed experiments, Acquired data A.C.: Performed experiments, Acquired data A.J.R.: Analyzed data

J.R.S.: Analyzed data

R.S.K.: Provided MLL AF4 B-ALL cell line R.D.D.: Designed research, Analyzed data

N.K.G.: Designed research, Analyzed data, Edited manuscript

D.S.R.: Designed research, Analyzed data, Wrote manuscript, Secured Funding, Project leader

