## Supplementary material for "A small molecule inhibitor of RNA-binding protein IGF2BP3 shows anti-leukemic activity": All Supplemental Information

**Supplementary Information for Jaiswal et al, 2025**

#### Detailed description of Methods

##### Cell lines and cell culture

All cell lines were maintained in standard conditions in incubator at 37 °C and 5% $CO_2$. Human B-ALL cell lines, RS4;11 (ATCC CRL-1873), NALM6 (ATCC CRL-3273), SEM (DMZ-ACC 546), PER785 , REH (ATCC CRL-8286 ), MOLM14 (DSMZ ACC 777), MV4;11 (ATCC CRL-9591), OCI-AML3 (DSMZ ACC 582 ) and KASUMI 1 (ATCC CRL-2724) were cultured as previously described (1). Immortalized MLL-Af4 transformed hematopoietic stem and progenitor cells derived from mouse bone marrow (MLL-Af4 Lin- cells) were cultured in IMDM with 15% FBS, supplemented with recombinant mouse stem cell factor (SCF, 100 ng/mL, Thermo Fisher), recombinant mouse Interleukin-6 (IL-6, 4 ng/mL, Thermo Fisher), recombinant human FMS like tyrosine kinase 3 ligand (FLT3-L, 50 ng/mL, Thermo Fisher) and mouse thrombopoietin (TPO,50 ng/mL, Thermo Fisher). OCI-AML8227 cells were a kind gift from Dr. John Dick and Dr. Eric Lechman (2). OCI-AML 8227 cells were cultured in a specialized media AMEM (Wisent Bio), BITS 9000 (Stem Cell Technologies), stem cell factor (SCF, 200 ng/mL, R&D systems), thrombopoietin (TPO, 25 ng/mL, R&D systems), FMS like tyrosine kinase 3 ligand (FLT3-L, 50 ng/mL, R&D systems), Interleukin-6 (IL-6, 20ng/mL, R&D systems), interleukin-3 (IL-3, 10 ng/mL, R&D systems), granulocyte Colony- Stimulating Factor (G-CSF, 10 ng/mL, R&D systems). OCI-AML8227 cells were cultured by plating 450000 cells/well of non-tissue culture treated 24-well plate (Eppendorf co.) and were passaged every 7 days. These are further listed in **Supp. Table 2**.

##### CRISPR/Cas9-mediated deletion of IGF2BP3 in cell lines

Human B-ALL cell lines SEM, RS4;11, and NALM6 were depleted for IGF2BP3 using lentiviral delivery of CRISPR/Cas9 components in a two-vector system and sgRNA sequence as previously described (1,3). Immortalized MLL-Af4 Lin- cells were initially isolated from bone marrow of Cas9-GFP mice and then transformed using retroviral transduction with MLL-Af4 retroviral supernatant, with four rounds of transduction with MLL-Af4 retroviral supernatant, followed by selection in G418 supplemented media at 400 μg/mL for 7 days, as previously described. Cells were then stably transduced with lentiviral supernatant containing sgRNA against *Igf2bp3* (I3Cr2) or non-targeting (NT-1), and sorted on GFP and mCherry positivity (1,3).

**Protein over-expression and purification**

IGF2BP3-GFP-Flag over-expression plasmid was constructed by fusing full length IGF2BP3 CDS with GFP CDS and cloned in pCDNA 3.0 vector. pCDNA3/IGF2BP3.GFP.Flag construct was transfected in 293T cells plated in 150 mm tissue culture dish. After 48 hours of transfection, cells were lysed in non-denaturing lysis buffer (50 mM Tris- HCl, pH 7.4, 300 mM NaCl, 5% glycerol, 0.1% NP40, 1X DNase, 10 units RNase/mL, 7.0 mg/50 mL of DTT, 1x Protease inhibitor) and incubated on ice for 1 hour. followed by 25 cycles of sonication with 10 seconds. 'on' and 20 seconds. 'off' pulse. Lysate was cleared by centrifugation at 12,000 x g for 15 minutes. Cleared lysates were loaded in the column filled with Anti-FLAG M2 resin (Sigma Aldrich) and passed multiple times to assist the binding. The bound protein fraction was eluted by adding 0.5 M Glycine HCl, pH 3.5 buffer. The eluent was concentrated using Vivaspin MwCO 50 column (Greiner Bio One). The concentrated protein was stored in 100 mM Tris-HCl buffer at pH 7.2. The purity and integrity of the protein was determined by mass spectroscopy and SDS-PAGE Commassie staining. The specificity of the purified protein was determined by western blotting and probing for IGF2BP3 using rabbit Anti-human IGF2BP3 antibody.

**Protein extraction and Western blot**

Cell lysates were made in non-denaturing cell lysis buffer and RIPA lysis buffer. Lysates were electrophoresed using SDS-PAGE using standard conditions (4). A comprehensive list of antibodies is provided in **Supp. Table 3**.

##### TR-FRET assay

TR-FRET assay was performed using the purified IGF2BP3-GFP fusion protein**.** Briefly, FRET assay was performed in assay buffer (50mM Tris-HCl, pH 7.4, 150 mM NaCl, DTT, 5% Glycerol, protease inhibitor (1x), RNase Inhibitor (10u/mL)). First, the purified protein was diluted at 50 nM concentration (2x) in the assay buffer and added to the plate. In the second step, 20 nM (2x) concentration of biotinylated M6A labelled RNA oligo (**Supp. Table 4**) was incubated with Streptavidin-Terbium in the assay buffer for 30 mins. on ice. After the incubation was over, the Streptavidin-Tb + RNA complex was added to the wells with purified proteins. The plate was incubated for 1 hr. at RT for binding and read on EnVision microplate reader (Perkin Elmer) with a dual PMT configuration using a 340 nm excitation and a dual emission mirror block with using a 495 nm emission filter for Terbium fluorescence (W1 channel) and a 525 nm emission filter for GFP fluorescence (W2 channel). The measurement details were as follows: 2000 cycles, 120 flashes, 200 us delay and 300 us total time window. Excitation and emission light were set to 100%. The assay was miniaturized to 10uL in 384-well plates and an in-house compound deck at UCLA MSSR was applied to the assay, with parameters as described in Results.

##### Data analysis of HTS assays

Values for W2, W1 and the ratio of W2 to W1 for each compound, DMSO and no IGF2BP3-GFP controls were imported into our Collaborative Drug Discovery platform for each plate and Z' for each plate determined. Z'<0.5 were flagged for repeats. Plates were normalized for Teribum (W1), GFP (W2) and W2/W1 ratios (Ratio), prospective hits were flagged for further analysis with a |Ratio Z| > 3 (Primary criterion). The secondary criterion was |W1 Z|<1 (where the mean and standard deviation were calculated for either each plate or for all the plates run on a given experiment). The “alternative secondary criterion” used the mean and standard deviation for all plates in a given experiment. Compounds with very strong inhibition in the W1 channel were treated as likely TRF-donor quenchers and de-prioritized. Compounds with very bright W2 signal were similarly treated as non-specifically fluorescent compounds. The residual structures of prospective hit compounds were exported for review by medicinal chemists and analysis through structure activity landscape index (SALI) plots(5). Following the identification of hits based on these parameters, we then designed a custom set of plates with compounds picked from the in house deck. The confirmatory TR-FRET screens were performed in triplicate with further analysis as described in the results section.

**Counter-Screen and Cell-based Assays.**

A custom set of plates was constructed from the hits that were identified from the initial TR-FRET based assays. Wild type and IGF2BP3-deleted SEM cell lines (3) were treated with compound, grown for 4 days under normal growth conditions, and assayed for cell growth in 384-well plates with CellTiter-Glo, a luminescence-based reagent described previously (1,3,6). Fold change from control (i.e., no compound treated wells) was plotted for SEM-WT versus SEM-I3KO. We utilized the cutoffs described in the results section to enable downstream analysis of potential lead compounds.

**Synthesis of I3IN-002**

Commercially available reagents were used as received. Methanol (MeOH) was purchased from Fischer Scientific. Deionized water was used. Dimethyl sulfoxide (DMSO) was purchased from Sigma-Aldrich. $K_2CO_3$ was purchased from Oakwood Chemical. 5-Methyl isatin **4** and $\alpha$-chloro amide **7** were purchased from Combi-Blocks. Thiosemicarbazide (**5**) was purchased from AK Scientific. Reaction temperatures above 23 °C were controlled using an IKAmag temperature modulator, and unless stated otherwise, performed at room temperature (approximately 23 °C). $^1$H-NMR spectra were recorded on Bruker spectrometers (at 500 MHz) and are reported relative to the residual solvent signal. Data for $^1$H-NMR spectra are reported as follows: chemical shift (d ppm), multiplicity, coupling constant (Hz), and integration. $^{13}$C-NMR spectra were recorded on Bruker spectrometers (at 125 MHz) and are reported relative to the residual solvent signal. Data for $^{13}$C-NMR spectra are reported as follows: chemical shift (d ppm) and, when necessary, multiplicity and coupling constant (Hz). $^{19}$F-NMR spectra were recorded on Bruker spectrometers (at 564 MHz) and are reported relative to the residual solvent signal. Data for $^{19}$F-NMR spectra are reported as follows: chemical shift (d ppm) multiplicity and integration. IR spectra were recorded on a Perkin-Elmer UATR Two FT-IR spectrometer and are reported in terms of frequency absorption ($cm^{-1}$). DART-MS spectra were collected on a Thermo Exactive Plus MSD (Thermo Scientific) equipped with an ID-CUBE ion source and a Vapur Interface (IonSense Inc.). Both the source and MSD were controlled by Excalibur software version 3.0. The analyte was spotted onto OpenSpot sampling cards (IonSense Inc.) using $CH_2Cl_2$ as the solvent. Ionization was accomplished using UHP He plasma with no additional ionization agents. The mass calibration was carried out using Pierce LTQ Velos ESI (+) and (–) Ion calibration solutions (Thermo Fisher Scientific). Retention times for liquid chromatography were recorded with a Shimadzu Nexera XR equipped with a PDA detector. Chromatography was conducted using a 0.5 mL/min flow rate at 40 °C and Shimadzu Nexcol C18 (1.8 x 50 mm, 2.1 μm) column. Two mobile phase solutions were used: solution A 0.1% formic acid in water and solution B was 0.1% formic acid in acetonitrile. The elution program consisted of a linear gradient starting at 95% A 0.5 min following injection to 5% A over 4 min. All the final compounds presented a purity of at least 95%, determined by HPLC and $^1$H NMR. Uncorrected melting points were measured using a Digimelt MPA160 melting point apparatus.

A round-bottom flask containing a magnetic stir bar was charged with $K_2CO_3$ (1.52 g, 11.0 mmol, 1.1 equiv), isatin **4** (1.75 g, 10.0 mmol, 1.0 equiv), thiosemicarbazide **5** (911 mg, 10.0 mmol, 1.0

equiv,) and MeOH (17 mL, 0.60 M). The reaction mixture was allowed to stir at 23 °C for 18 h. At this point, the reaction mixture was acidified by dropwise addition of acetic acid over roughly 5 min (5 mL). The resulting mixture was stirred for 1 hour. After 1 h, the mixture was vacuum filtered over a fritted funnel, and the filter cake containing the product was washed with $H_2O$ (100 mL), followed by $Et_2O$ (50 mL) to afford (E)-2-(5-ethyl-2-oxoindolin-3-ylidene) hydrazine-1-carbothioamide (2.10 g, crude mass) as a yellow-orange solid. This was used directly in the subsequent reaction without further purification.

To a round-bottom flask containing a magnetic stir bar, (E)-2-(5-ethyl-2-oxoindolin-3-ylidene) hydrazine-1-carbothioamide (2.10 g, 8.46 mmol, 1.0 equiv.) and $K_2CO_3$ (1.17 g, 8.46 mmol, 1.0 equiv) were added, and dissolved in $H_2O$ (100 mL, 0.08 M). The reaction vessel was fitted with a reflux condenser and the reaction mixture was stirred at 100 °C for 16 h. At this point, the crude mixture was allowed to cool to 23 °C. It was acidified by the dropwise addition of acetic acid over roughly 5 min (10 mL) and then stirred for an additional hour. The mixture was vacuum filtered over a fritted funnel, and the filter cake containing the product was washed with $H_2O$ (100 mL), followed by $Et_2O$ (50 mL) to afford triazinoindolothione **6** (1.47 g, 65% yield over two steps, >95% purity) as a yellow-orange solid.

To a 1-dram vial charged with a magnetic stir bar was added triazinoindolothione **6** (50 mg, 0.22 mmol, 1.0 equiv), $\alpha$-chloroacetamide **7**(52 mg, 0.22 mmol, 1.0 equiv), $K_2CO_3$ (60 mg, 0.43 mmol, 2.0 equiv), and DMSO (1.1 mL, 0.20 molar). The reaction mixture was allowed to stir at 23 °C for 16 h. The reaction mixture was transferred to a vial containing $H_2O$ (5 mL), which led to the formation of a precipitate. The mixture was sonicated for 5 minutes to afford the crude product as a suspension in water. This mixture was then heated to reflux for ~ 5 seconds to effect partial dissolution, and was then allowed to cool to 23 °C under ambient conditions. The product precipitated and was collected by vacuum filtration over a paper filter. The filter cake was successively washed with $H_2O$ (5 x 1 mL), MeOH (1 x 0.5 mL), and $Et_2O$ (2 x 1 mL) to give **I3IN-002** (42 mg, 45% yield, >95% purity) as a tan solid. **I3IN-002**: Mp: 253 °C; $^1$H NMR (500 MHz,

$(CD_3)_2SO$): δ 12.54 (s, 1H), 9.90 (s, 1H), 8.14 (s, 1H), 7.72 (d, *J* = 7.8 Hz, 1H), 7.68 (t, *J* = 7.8 Hz, 1H), 7.60 (d, *J* = 8.0 Hz, 1H), 7.56 (dd, *J* = 8.3, 1.4 Hz, 1H), 7.51 (d, *J* = 8.3 Hz, 1H), 7.44 (t, *J* = 7.6 Hz, 1H), 4.27 (s, 2H), 2.80 (q, *J* = 7.5 Hz, 2H), 1.27 (t, *J* = 7.6 Hz, 3H); $^{13}C$ NMR (125 MHz, $(CD_3)_2SO$): δ 167.3, 165.9, 146.6, 141.3, 138.7, 138.4, 135.2, 133.1, 131.2, 129.3, 126.6, 126.3 (q, *J* = 5.0 Hz), 124.0 (q, *J* = 29.6 Hz), 123.5 (q, *J* = 273.4 Hz), 120.1, 117.6, 112.6, 34.5, 28.0, 16.2; $^{19}F$ NMR (564 MHz, $(CD_3)_2SO$): δ –59.4 (s, 3H); IR (Film): 3263, 2961, 1662, 1536, 1095 $cm^{-1}$; HRMS-APCI (*m/z)* $[M + H]^+$ calcd for $C_{20}H_{17}F_3N_5OS^+$, 432.11004; found 432.11002; HPLC purity 97 %; $R_t$ = 4.20 min.

**In vitro assays for cell viability, cell cycle apoptosis, and proliferation, with small molecule treatment**

Cell viability assays were performed using a luminescent assay based on ATP quantitation (CellTiterGlo, Promega, catalog G7571), per manufacturer's protocol. 1500 cells/well (for 384 well format) and 5000 cells/well (for 96 well format) were plated and incubated with different concentrations of small molecule for four days at 37°C in a 5% $CO_2$ incubator, followed by endpoint assay. For $IC_{50}$ determinations, small molecule was added at concentrations from 50 μM-0.0004 μM, with 2-fold dilutions. For cell cycle analysis, SEM and RS4;11 cells were treated with 5 μM concentration of I3IN-002 and DMSO for 48 hrs. After 48 hrs. cells were harvested, washed with PBS, and fixed in pre-chilled 70% ethanol overnight. Fixed cells were washed with PBS and 350 µL Propidium Iodide staining solution (20 µg/mL of propidium Iodide and 200 µg/mL of DNase free RNase) was added to the cells and incubated for 1 hr. at RT. Stained cells were run on the flow cytometer and analyzed using FlowJo software V10.

**Apoptosis**

Apoptosis assays were performed using luminescence-based Caspase Glo assay (Promega). Briefly, SEM and RS4;11 cells were treated with 5 µM concentration of I3IN-002 and DMSO for 48 hrs. Post treatment, 100 µL of Caspase Glo reagent was added to each well (1:1) and incubated for 1 hr. at 37 degree Celsius. Plate was read on Varioscan lux microplate reader from Thermo Scientific. The increase in luminescence in I3IN-002 treated cells compared to DMSO corresponds to an increase in Caspase 3/7 activity. An alternate measurement of apoptosis was performed on the compound treated cells using Annexin V staining as described previously(1).

**Colony formation assay**

Colony formation assay with serial replating was performed using murine Lin-MLL-Af4 cells(3). Briefly, 6000 cells were resuspended in 4 mL of Methocult media (M3434, Stem Cell Technologies) with DMSO/I3IN-002. Then, 1.1 mL of the cell suspension was plated in each 35 mm petri dish and incubated for 8 days. After colony counting, colonies were pooled by adding PBS at room temperature, creating a single-cell suspension by gently mixing with a micropipette, and centrifuging at 1200 rpm for 5 minutes. The pellet was resuspended in PBS, and viable cells were counted using Trypan blue. For serial replating, 1600 cells were seeded again following the previously mentioned steps with DMSO and I3IN-002.

**Animal Experiments**

For in vivo studies, C57BL/6J, B6.SJL-*Ptprc*$^{a}$ *Pepc*$^{b,}$/BoyJ (B6 CD45.1), and B6J.129(Cg)-Gt(ROSA)26Sor$^{tm1.1(CAG-cas9_{*},-EGFP)Fezh/J}$ (Cas9-GFP, BL/6J) were procured from The Jackson Laboratory. Primary murine leukemia cells (WT and Igf2bp3 KO) as previously described (3) were transplanted into busulfan-conditioned recipients. One week following transplantation, mice were injected with vehicle or I3IN-002, three times a week intraperitoneally, at a dose of 25 mg/kg, for three weeks, with an endpoint at four weeks post-transplantation. The peripheral blood engraftment of the leukemic cells was checked by FACS at week 2 and week 4. Once the peripheral blood engraftment reached >20% at 4 weeks, the experiment was terminated and tissues were harvested to be analyzed by FACS, histology and RT-qPCR. All of the animal experiments received Institutional Animal Research Committee approval at UCLA.

**Toxicity Study protocol**

To study the safety of the IGF2BP3 inhibitor I3IN-002, we performed a toxicity study on 8-10 week-old C57BL6/J mice. The mice were injected with 50 µL of vehicle and I3IN-002 at a dose of 25 mg/kg body weight, three times a week for a period of four weeks. During the course of the study, body weight was measured every week and CBC analysis performed every two weeks. The experiment was terminated at the end of four weeks, and the mice were sacrificed to collect cells from the bone marrow and spleen to study the effect of the inhibitor on the cells of the hematopoietic lineage. Peripheral blood was collected from the terminal bleed through heart puncture to isolate serum for serum chemistry analysis. Tissues such as the liver, kidneys, lungs, thymus, heart, and brain were harvested for histological examination. All of the studies on animals were approved by the Institutional Biosafety Committee at UCLA.

**RNA immunoprecipitation and qPCR assays.**

RNA immunoprecipitation assay was performed on SEM cells treated with 5 µM concentration of I3IN-002 and DMSO for 48 hours. After incubation, cells were lysed in a non-denaturing lysis buffer supplemented with protease inhibitor and RNase inhibitor. Lysate was cleared at 12000 rpm for 15 minutes. and RIP was performed on the cleared lysate by adding 5 µg of IGF2BP3 antibody/ IgG and 50 µL of Protein G agarose beads and incubated overnight at 4 degree Celsius. RIP and qRT-PCR were performed using protocol described previously.

**Cellular thermal shift assay (CETSA)**

A cellular thermal shift assay was performed on SEM cells treated with the test compound, based on published protocols(7). Briefly, 20 x$10^6$ cells were treated with 25 µM concentration of test compound in 40 mL of complete growth media and incubated for 24 hour. at 37 ˚C in $CO_2$ incubator, DMSO was used as vehicle control. After incubation, cells were spun down and washed with PBS to remove any trace of test compounds. Cells were further re-suspended in 1 mL of PBS with 1x protease inhibitor cocktail and 100 uL of cell suspension was distributed in individual PCR tubes. Cells were subjected to thermal denaturation at different temperature ranging from 38 ˚C – 52 ˚C on a thermal cycler for 3 minutes. Thermal treated cells were lysed by flash freeze and thaw cycle in liquid Nitrogen and cleared by centrifugation at 20,000xg for 20 minutes. at 4 °C. Lysates were boiled in sample buffer and subjected to western blotting.

**Drug Affinity responsive target stability (DARTS) assay**

DARTS assay (8) was performed to study direct binding of I3IN-002 with IGF2BP3 protein. Briefly, 5x$10^7$ SEM cells were lysed in 1 mL of m-PER buffer (Thermo scientific) and incubated on ice for 10 minutes. After incubation, lysate was centrifuged at 17,000xg for 10 min at $4^0$C. The supernatant was transferred to a fresh tube and mixed with one-tenth of the 10x TNC buffer (500 mM Tris-HCl, pH 8.0, 500 mM NaCl and 100 mM $CaCl_2$), followed by protein estimation using the BCA assay. Approximately 50 µg of lysate was incubated with different concentrations of I3IN-002 (DMSO, 25, 50,100, 250 and 500 µM) for 1 hour, followed by pronase (Roche, 10,165,921,001) treatment (1:500) for 30 minutes at room temperature. The pronase activity was quenched by adding a protease inhibitor cocktail, and the sample was perpared using SDS loading buffer and boiled for 10 min at $90^0$C. The samples were then used to run a western blotting assay, probing for IGF2BP3. Actin beta was used as a control.

**Thermal Shift Assay (TSA)**

Thermal shift assay (9) was performed using both full-length IGF2BP3 and RRM1-2 domain of IGF2BP3 purified protein. Briefly, 10 μM of purified protein was incubated with 100 μM of I3IN-002 for 30 minutes at room temperature. SYPRO dye (5000x) diluted in Bis-Tris buffer, pH 6.5 was added to the reaction mix. The reaction was run on a quantitative PCR machine (QuantStudio 3) using melt curve protocol.

**RNA isolation and qPCR**

Previous protocols were adapted for RT-qPCR as standard procedures (4). A full list of RT-qPCR primers are presented in **Supp. Table 5***.* For normalization to housekeeping genes, we used RT-qPCR primers for human 18S (or actin) and mouse L32.

**RNA Sequencing, Bioinformatics Methods & Data Analysis**

Total RNA was extracted from cell pellets using Qiazol (Qiagen) per manufacturer's protocol with the following modification to include an additional RNA ethanol wash step: after the total RNA was solubilized in dd$H_2O$, one overnight ethanol precipitation step was included for further purification. Libraries for RNA-Seq were prepared with KAPA mRNA-Seq Hyper Prep Kit. The workflow consisted of mRNA enrichment and fragmentation, first strand cDNA synthesis using random priming followed by second strand synthesis converting cDNA:RNA hybrid to double-stranded cDNA (dscDNA), and incorporating dUTP into the second cDNA strand. cDNA generation was followed by end repair to generate blunt ends, A-tailing, adaptor ligation and PCR amplification. Different adaptors were used for multiplexing samples in one lane. Sequencing was performed on Illumina HiSeq3000 for a SE 1x50 run. Data quality check was done on Illumina SAV. Demultiplexing was performed with Illumina Bcl2fastq v2.19.1.403 software. The reads were mapped by STAR 2.7.9a(10) and read counts per gene were quantified using the human genome GRCh38. In Partek Flow, read counts were normalized by CPM +1.0E-4. All results of differential gene expression analysis utilized the statistical analysis tool, DESeq2(11). P-value ($p<0.05$), FDR

(<0.05), and fold change (FC>2-fold) filters were applied for differentially expressed gene lists prior to downstream analysis. Multiple testing correction was performed using the Benjamini–Hochberg method. Significant differentially expressed genes have adjusted *P* value =< 0.05 and absolute log2FC >= 1. Enrichment analyses were completed with Metascape (12).

**Statistical analysis**

Data shown represents mean ± SD for continuous numerical data. Two-tailed student's *t* tests or one-way ANOVA followed by Bonferroni's multiple comparisons test were performed using GraphPad Prism software and conducted as described in the figure legends. Survival analyses were performed using Kaplan-Meier method with comparisons made using log-rank tests, followed by Bonferroni's correction for multiple comparisons. A *P* value less than 0.05 was considered significant.

**Additional reagents.** Additional reagents and kits are listed in **Supp. Table 5**.

**Supplemental Figure Legends.**

**Supplemental Figure 1, related to Figure 1. Configuration and standardization of TR-FRET assay.**

a. Schematic representation of constructs used in protein purification for TR-FRET.

b. Photomicrograph of GFP fluorescence following transfection of plasmid. Scale bar, 50 micron.

c. Western blot analysis for IGF2BP3 in elutions from anti-Flag column.

d. Coomassie stained polyacrylamide gel showing I3-GFP purified from the column.

e-g. RNA immunoprecipitation using purified IGF2BP3 protein (confirmed by Western Blot), (e) of *CDK6* (f) and *MYC* (g) mRNAs, assessed by RT-qPCR.

h. GFP Fluorescence (arbitrary units) as a function of RNA oligonucleotide concentration.

i-j. GFP fluorescence and Z' values in three different experiments performed on three different days (E1-E3; n=110 (replicates); assay volume= 10 µL)

k. Comparison of GFP fluorescence in 100 µL and 10 µL assay volume**.**

**Supplemental Figure 2. High throughput screening assay, related to Figure 1.**

a. Plate layout and example plate with hit (right). Please see materials and methods for further details.

b. Plot of plate standard deviations across 600+ plates used for the assay.

c. Number of compounds identified as hits by different cut-off criteria in preliminary screening experiment. Based on established literature and these results a cut-off of $|z|>3$ was chosen.

d. To confirm hits identified in the full-scale screen, several replicate experiments were carried out to assess changes in fluorescence ratios. 417 high-probability hits were re-pinned into two 384-well plates and were subjected to the same TR-FRET assay as before. An individual compound was defined as being a "hit" in the validation screen when the fluorescence ratio deviated from negative controls by greater than three standard deviations. Next, we defined a

Tier 1 hit compound as positive in both experimental replicates; Tier 2 hit compound as positive in 1 out of 2 experimental replicates, and Tier 3 compounds were "non-hits" in the validation experiment based on their not being positive in either replicate. The relative frequency of Tier 1, Tier 2 and Tier 3 compounds amongst all "primary hits" versus amongst compounds with "Core structure" are plotted here.

e. Overall strategy of TR-FRET screen, validation, and counter-screen.

**Supplemental Figure 3. Characterization of anti-leukemic effects of I3IN-002, related to Figure 3.**

a-b. FACS-based cell cycle analysis following treatment with I3IN-002 versus control, with histograms of propidium iodide staining in SEM (a) and RS4;11 cells (b).

c. Quantification of percentage of RS4;11 cells at each stage of cell cycle with 3 replicates.

d-e. G2-M transition genes CDK1 and CCNB2 downregulated in I3IN-002 treatment.

f. Measurement of apoptosis induction, by Caspase 3/7 activity assay in RS4;11 following treatment with I3IN-002. Data represents mean and standard error of mean with 8 replicates.

g-i. FACS scatter plot of propidium iodide/Annexin V co-staining of RS4;11 cells (left) with quantitation of total Annexin V positivity in RS4;11 and SEM cells following treatment with I3IN-002. Data represents mean and standard error of mean with 3 replicates.

j. Decreased colony counts upon serial replating in different treatment groups.

**Supplemental Figure 4. Further characterization of in vivo phenotypes.**

a. Western blot showing Igf2bp3 expression in NT and Igf2bp3 knockout Lin- MLL- Af4 cells.

b. FACS histograms of CD45.1/CD45.2 congenic marker staining for measurement of leukemia engraftment.

c. FACS histograms of CD45.2/CD11b staining for measurement of leukemia engraftment.

d-e. Absolute counts of CD11b and CD34+ Sca1- cells in bone marrow compartments.

f. Histologic sections of lung and kidney comparing the morphology of tissues following engraftment of leukemia and treatment with vehicle, I3IN-002, or genetic deletion of IGF2BP3. Arrows show areas of leukemic infiltration. Hematoxylin and eosin staining, original magnification, 200X.

**Supplemental Figure 5. Characterization of on-target activity of IGF2BP3, related to Figure 5.**

a-b. RT-qPCR analysis for *MYC* and *HOXA9* from RNA immunoprecipitation sample.

c. Drug affinity responsive target stability show I3IN-002 protects IGF2BP3 from Pronase activity.

d. IGF2BP3 expression in NT and IGF2BP3 KO SEM cells.

e-f. RT-qPCR of additional genes identified from differential expression analyses of RNA-sequencing experiments (*KDM3A* and *TGFBR2*, respectively).

g-l. RT-qPCR analyses of RNA isolated from mouse spleen following in vivo treatment described in **Fig. 4**. Three representative mice from each group were selected for the RT-qPCR analyses.

Supplementary Figure 1.

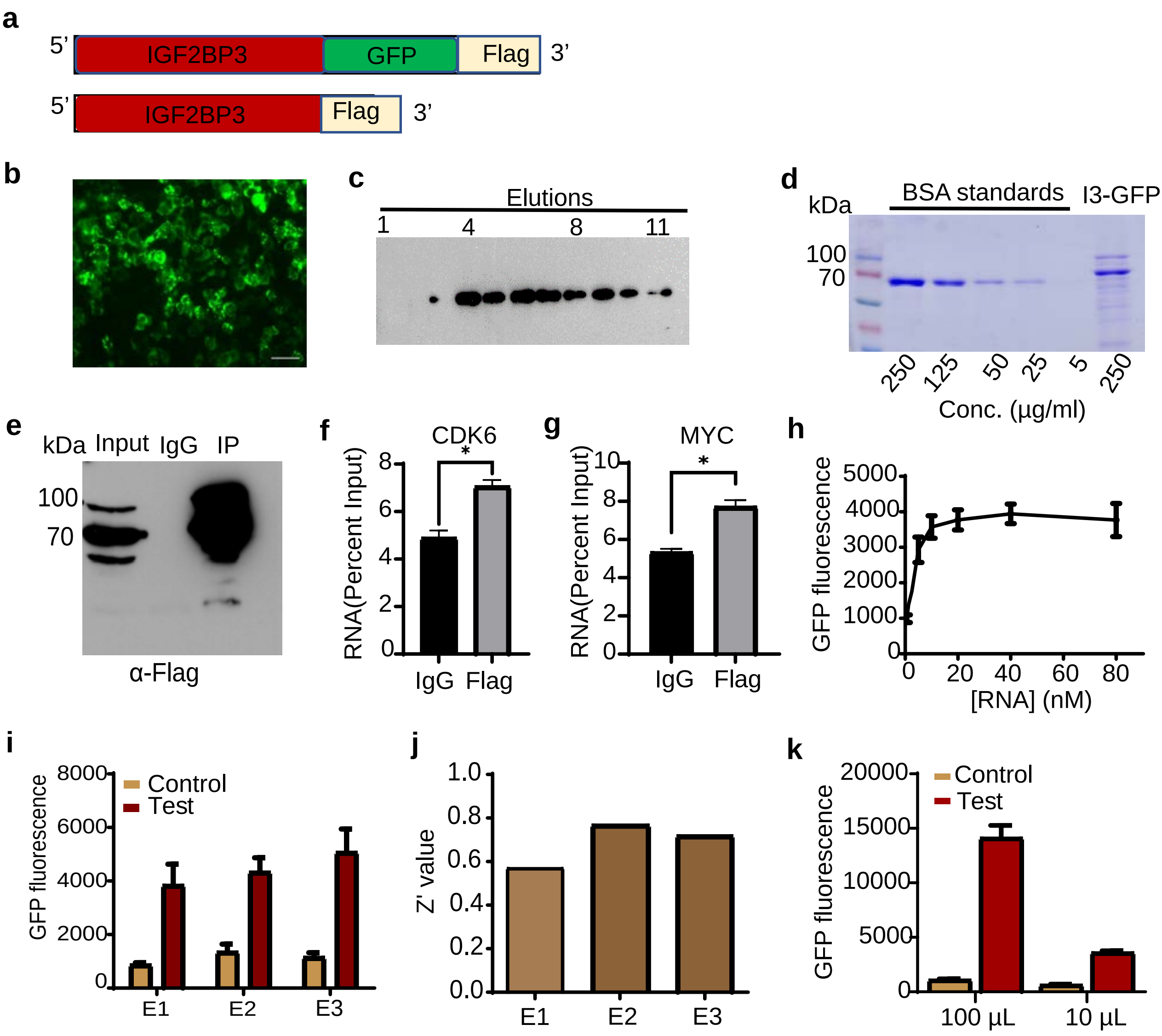

#### Supplementary Figure 2.

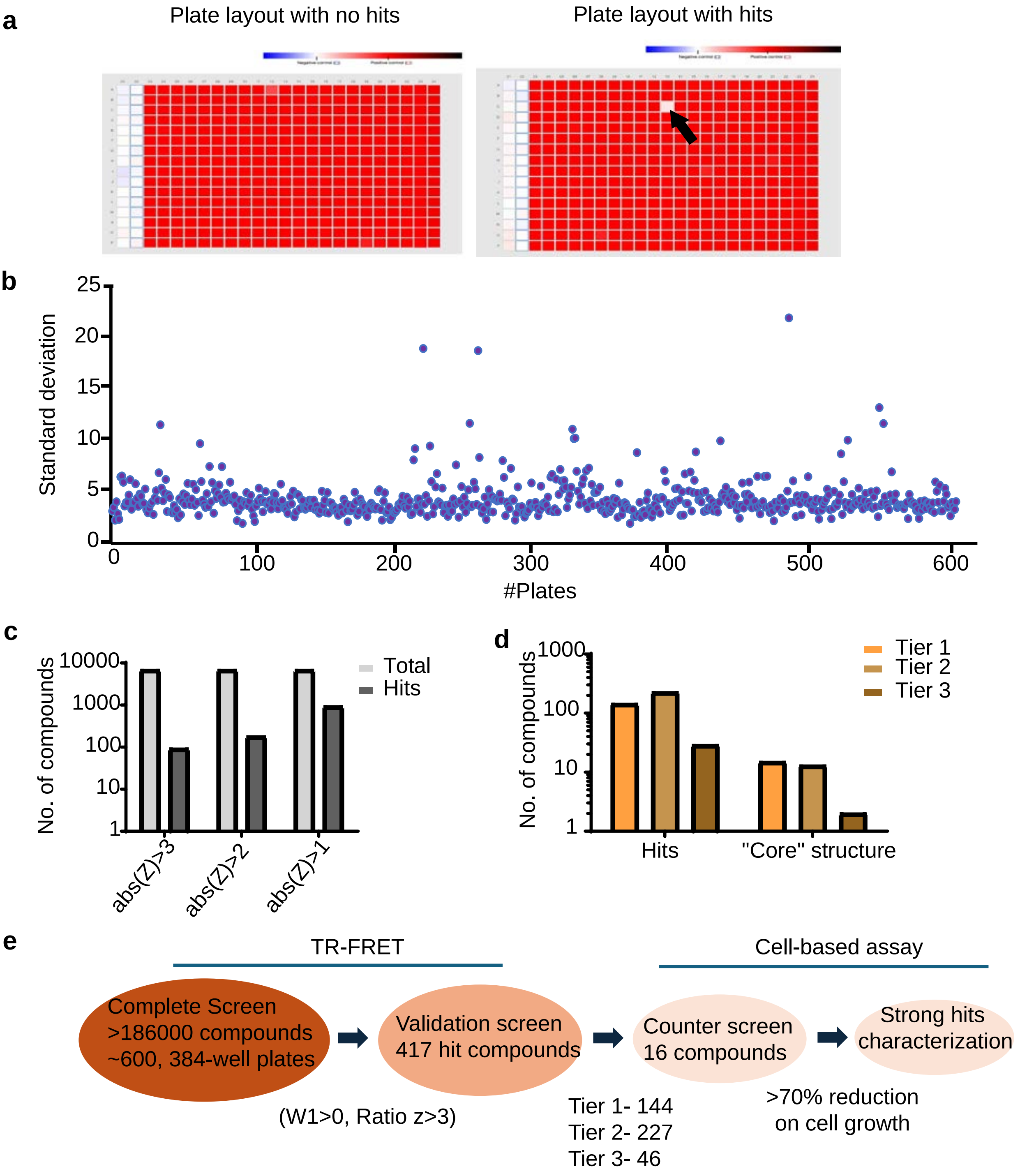

#### Supplementary Figure 3.

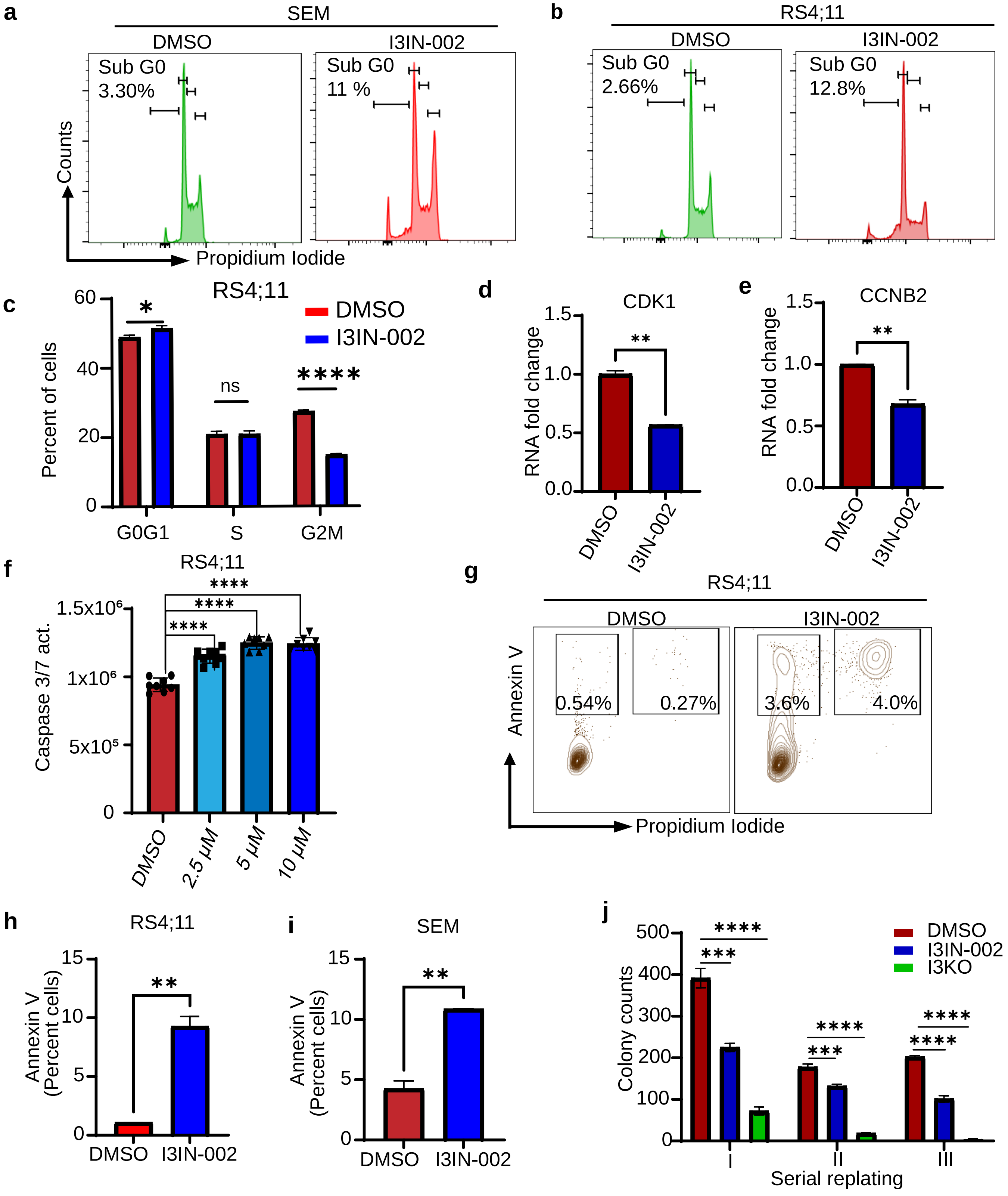

### Supplementary Figure 4.

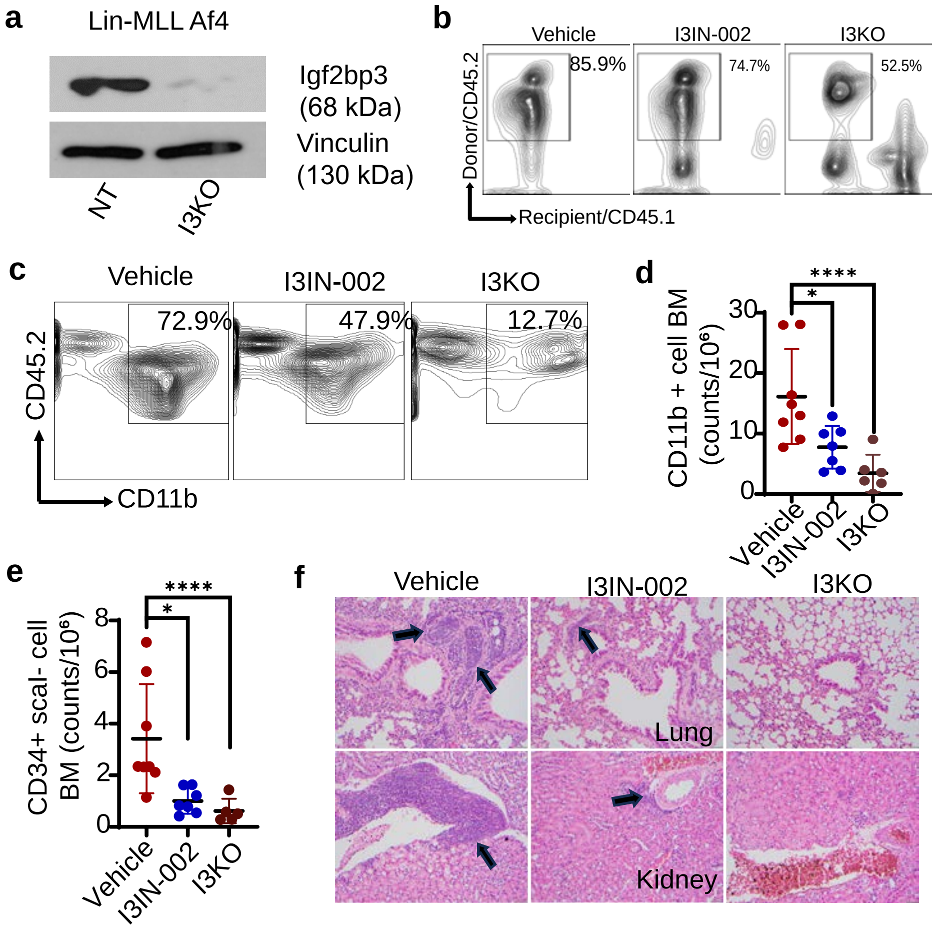

#### Supplementary Figure 5.

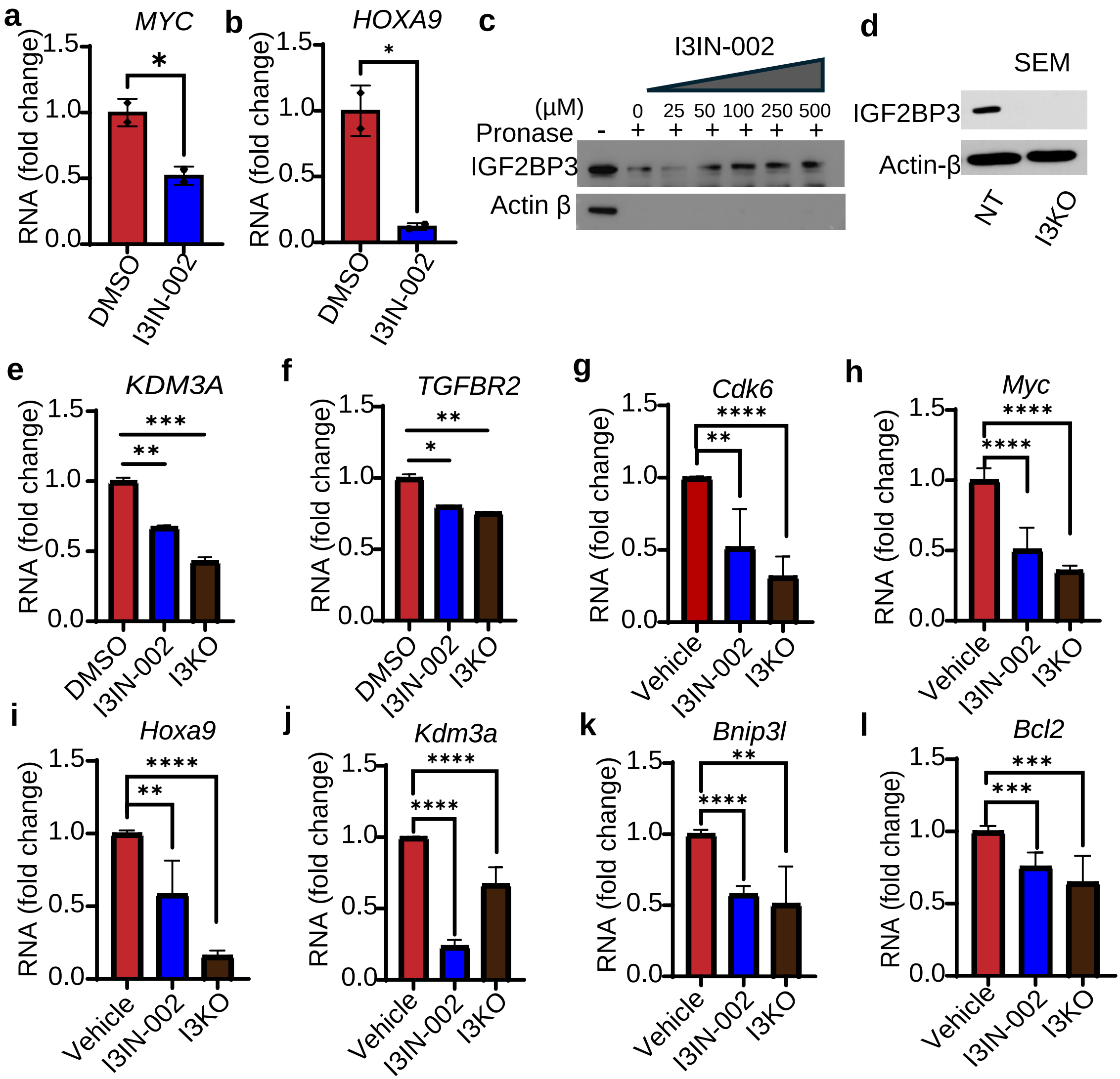

**Supplemental Table 1. Reagents used in cell culture experiments.**

| Cell lines and media components | | |
|---|---|---|
| RS4;11 | ATCC | CRL-1873 |
| NALM6 | ATCC | CRL-3273 |
| SEM | DSMZ | ACC 546 |
| IMDM | Gibco | 12440046 |
| RPMI 1640 | Gibco | 11875119 |
| FBS | Gibco | 26140079 |
| mSCF | Invitrogen | RP87738 |
| IL-6 | Gibco | PHC0065 |
| hFLT-3 | Gibco | PHC9415 |
| mTPO | Invitrogen | RP87753 |
| AMEM | Wisent | 310-010 CL |
| BITS 9000 | Stemcell Technologies | 09500 |
| R&D SCF | R&D Systems | 11010SC010 |
| R&D TPO | R&D Systems | 288TP005 |
| R&D FLT3-L | R&D Systems | 308FK025 |
| R&D IL-6 | R&D Systems | 206IL050 |
| R&D IL-3 | R&D Systems | 203IL010 |
| R&D G-CSF | R&D Systems | 214CS005 |

**Supplemental Table 2. Antibodies used in FACS, western blot and immunoprecipitation assays.**

| Antibodies | | |
|---|---|---|
| Biolegend | 105826 | APC/Cyanine7 anti-mouse CD117 (c-kit) |
| Biolegend | 118103 | Biotin anti-mouse TCR γ/δ Antibody |
| Biolegend | 312225 | Brilliant Violet 711™ anti-human CD10 Antibody |
| Biolegend | 157211 | Pacific Blue™ anti-mouse CD45 Antibody |
| Biolegend | 101318 | PE/Cyanine7 anti-mouse CD16/32 Antibody |
| Biolegend | 108124 | PerCP/Cyanine5.5 anti-mouse Ly-6A/E (Sca-1) |
| Biolegend | 343513 | APC/Cyanine7 anti-human CD34 Antibody |
| Biolegend | 303521 | PerCP/Cyanine5.5 anti-human CD38 Antibody, |
| Biolegend | 135120 | Brilliant Violet 605™ anti-mouse CD117 (c-kit) Antibody |
| Biolegend | 109815 | Alexa Fluor® 488 anti-mouse CD45.2 Antibody |
| Biolegend | 101215 | PE/Cyanine7 anti-mouse/human CD11b Antibody |
| BD Bioscience | 563973 | Annexin V BV421 |
| BD Bioscience | 556463 | Propidium Iodide solution |
| Proteintech | 67447-1-Ig | cmyc monoclonal antibody |
| Fisher | 50-173-2168 | HOXA9 Rabbit anti-Human |
| Fisher | PIPA527978 | CDK6 Polyclonal Antibody |
| Santa Cruz Biotech | sc-2357 | mouse anti-rabbit IgG-HRP |
| Proteintech | 146421AP150UL | rabbit anti-human igf2bp3 |
| MBL | RN009P | MBL igf2bp3 rabbit anti-human/mouse |
| Sigma Aldrich | A5441 | Actin beta Ab |
| Sigma Aldrich | F1804 | Anti-FLAG Ab |
| Proteintech | 674471IG150UL | Myc anti-human |
| Invitrogen | PA527978 | CDK6 anti-human |
| Proteintech | 185011AP150UL | HOXA9 rabbit anti-human |
| Invitrogen | PA527094 | BCL-2 mouse anti-human |

**Supplemental Table 3. RNA Sequences used in IGF2BP3-RNA time resolved Forester resonance energy transfer assay.**

| RNA Oligo for FRET | Sequence |
|---|---|
| Unmethylated | 5’biotin- CGU CUC GGA CUC GGA CUG CU-3’ |
| Methylated | 5’biotin- CGU CUC GG($m^6$A) CUC GG($m^6$A) CUG CU-3’ |

**Supplemental Table 4. Quantitative PCR primers for genes assayed.**

| Gene name | Primer Sequence |
|---|---|
| mRPS32-F | AAGCGAAACTGGCGGAAAC |
| mRPS32-R | TAACCGATGTTGGGCATCAG |
| TGFBR2-F | AAGATGACCGCTCTGACATCA |
| TGFBR2-R | CTTATAGACCTCAGCAAAGCGAC |
| BNIP3L-F | CAGCAATAATGGGAACGGGG |
| BNIP3L-R | ATCTTGTGGTGTCTGCGAGC |
| P4HA1_F | AGGACTGCTTTGAGTTGGGC |
| P4HA1_R | TCTCGCCTTCATCCAGTTGC |
| LSP1 F | TTGAGTCTGAGCAAGGAGGG |
| LSP1 R | CTGGGCTGCTGACATTTCTG |
| P4HA1 F | ATGACCCCTCGGAGACAGAA |
| P4HA1 R | GCCTCAGCCTTGGTTTTGC |
| MYC F | CCACAGCAAACCTCCTCACAG |
| MYC R | GCAGGATAGTCCTTCCGAGTG |
| CDK6 F | GCTGACCAGCAGTACGAATG |
| CDK6 R | GCACACATCAAACAACCTGACC |
| HOXA9 F | TGGACAGACTTAAATGCCCGC |
| HOXA9 R | TGAACCTATGATTGTAAGGAGCTG |
| BCL2 F | CGG TGG GGT CAT GTG TGT G |
| BCL2 R | CGG TTC AGG TAC TCA GTC ATC C |
| KDM3A F | CAGGAGCCACAGTAGGAGAC |
| KDM3A R | AGTTTGCCATCTCGCCTTGT |
| RPS18 F | GAGGATGAGGTGGAACGTGT |
| RPS18 R | GGACCTGGCTGTATTTTCCA |
| Cdk6 F | TCTCACAGAGTAGTGCATCGT |
| Cdk6 R | CGAGGTAAGGGCCATCTGAAAA |
| Bcl2 F | GTC GCT ACC GTC GTG ACT TC |
| Bcl 2 R | CAG ACA TGC ACC TAC CCA GC |
| Hoxa9 F | AAAACACCAGACGCTGGAAC |
| Hoxa9 R | TCTTTTGCTCGGTCCTTGTT |
| Myc F | ATG CCC CTC AAC GTG AAC TTC |
| Myc R | CGC AAC ATA GGA TGG AGA GCA |
| Kdm3a F | TCGGAGACTTCTGGGATGGA |
| Kdm3a R | TTCAGTTTGCCATCTCGCCT |
| Bnip3l F | AGACCCGAAAACATCCCACC |
| Bnip3l R | CAGAAGGTGTGCTCAGTCGT |

**Supplemental Table 5. Source of kits and reagents used as described in methods.**

| Kits and Reagents | | |
|---|---|---|
| Promega | G8091 | Caspase-Glo(R) 3/7 Assay, 10ml |
| Promega | G7572 | CellTiter-Glo(R) Luminescent Cell Viability, 100m |
| Corning | 21030CV | PBS |
| Thermo Scientific | EN0531 | DNase free Rnase A |
| Invitrogen | NP0007 | Invitrogen™ NuPAGE™ LDS Sample Buffer (4X) |
| Invitrogen | NP0002 | Invitrogen™ NuPAGE™ MES SDS Running Buffer (20X) |
| VWR | 101414-278 | PerfeCTa® SYBR® Green FastMix®, ROX™ |
| VWR | 101414-106 | qScript cDNA SuperMix, 5X, 100 rnxs |
| Thermo Scientific | 78438 | Protease inhibitor cocktail |
| Thermo Scientific | EO0381 | RNase inhibitor |
| Sigma Aldrich | A2220 | A2220 - ANTI-FLAG ® M2 Affinity Gel |
| Qiagen | 12162 | QIAGEN Plasmid Maxi Kit (10) |
| Qiagen | 217004 | miRNeasy Mini Kit (50) |
| Sigma Aldrich | B2635-10G | Busulfan |
| Sigma Aldrich | PPB019 | Bis-Tris-Bicine-EDTA |
| Sigma Aldrich | 11719408001 | Protein A Agarose, 2 mL |
| StemCell Tech | 3434 | MethoCult |
